# Preserved functions with profound morphological reorganization in human organotypic cultures

**DOI:** 10.64898/2026.05.08.723812

**Authors:** Estilla Zsófia Tóth, Rebeka Stelcz, Réka Bod, Ábel Petik, Eszter Juhász, Kinga Tóth, Liza Szalai, Nour Essam, Fanni Somogyi, Beatrix Kovács, Amirmahdi Mojtahedzadeh, Attila György Bagó, Loránd Erőss, Péter Orbay, Johanna Petra Szabó, Dániel Fabó, Boglárka Hajnal, Bence Rácz, Daniel Hillier, István Ulbert, Lucia Wittner

## Abstract

Human organotypic slice cultures provide experimental access to adult human neuronal circuits ex vivo, yet it remains unclear whether these networks preserve function or undergo fundamental reorganization following the profound perturbation of slice preparation.

Here, we combined extracellular population and single-unit electrophysiology, longitudinal calcium imaging, and quantitative histology to track the changes of human cortical slice cultures over several weeks in vitro. Early phases were marked by pronounced variability and instability, with reduced firing rates, increased burst propensity in principal cells, elevated discharge irregularity, and heterogeneous recruitment during population activity. These functional changes coincided with substantial structural remodeling, including considerable neuronal loss, disruption of laminar architecture, reactive gliosis, and selective vulnerability of inhibitory interneurons.

Strikingly, despite this progressive structural degradation, neuronal activity did not diverge but instead converged. By the fourth week in culture, electrophysiological properties, cell-type-specific firing patterns, and population-event recruitment became stable and highly consistent across patients. Calcium imaging revealed persistent, spatially confined regions of synchronous activity, indicating the preservation of structured network dynamics. These events remained within physiological regimes and lacked features of epileptiform discharges.

Thus, human neuronal circuits exhibit a robust capacity for self-organization, transitioning from heterogeneous, injury-driven dynamics to stable and homogeneous functional states. This dissociation between structural deterioration and functional convergence establishes human organotypic slice cultures as a reproducible and translationally relevant platform for studying human brain network dynamics ex vivo.

## Introduction

Understanding how human neuronal circuits respond to injury while preserving functional output remains a central challenge in neuroscience. In vivo human studies provide limited access to cellular and circuit-level mechanisms, whereas reductionist in vitro systems lack the architectural and cellular complexity of intact brain networks. Organotypic slice cultures bridge this gap by preserving elements of native cytoarchitecture, local connectivity, and cell-type diversity while enabling long-term experimental access to neuronal circuits^1^.

A critical distinction, however, lies between rodent and human organotypic models. Rodent slice cultures are typically derived from embryonic or early postnatal tissue, in which networks are still undergoing developmental maturation^2^. In contrast, human organotypic slices are prepared from adult surgical resections, often originating from epileptic or tumour-associated cortex, and therefore represent fully developed and functionally specialized circuits with prior physiological and pathological history^3,4^. This fundamental difference raises an important unresolved question: to what extent do adult human neuronal networks retain their intrinsic functional organization after the profound perturbation of slice preparation?

Studies in rodent organotypic cultures have established that slice preparation induces a cascade of network-level changes, including dendritic and axonal transection, synaptic reorganization, and homeostatic plasticity, which together reshape neuronal activity over days to weeks in vitro^5^. These processes are typically associated with dynamic changes in firing rates, burst structure, and synchronization, reflecting the transition from an acutely deafferented state to a re-equilibrated network. In human tissue, recent work has demonstrated the persistence of spontaneous and evoked activity in organotypic cultures^6–8^, as well as the maintenance of certain cell-type-specific physiological properties^4^. However, a comprehensive understanding of how single-neuron activity, population dynamics, and morphological properties co-evolve over time in human slice cultures remains lacking.

Another unresolved issue concerns variability. Human surgical samples are inherently heterogeneous, reflecting differences in age, pathology, anatomical origin, and prior treatment. This variability poses a challenge for reproducibility but also offers an opportunity to assess whether intrinsic circuit mechanisms drive convergence toward stable functional states^9^. In rodent systems, network activity is strongly influenced by developmental stage and genetic background, often resulting in substantial variability across preparations^10^. Whether similar or distinct principles govern the evolution of human neuronal networks in vitro is not well understood.

To address these questions, we performed a multimodal analysis of human organotypic slice cultures, combining extracellular population and single-unit electrophysiology, longitudinal calcium imaging, and quantitative histological assessment across extended culture periods. This approach enables simultaneous investigation of neuronal activity at multiple scales: from spike timing and cell-type-specific firing patterns to mesoscale population dynamics and underlying morphological organization. We hypothesized that the substantial structural perturbation predicted from the culturing methodology will profoundly affect the functional output. In contrast, we found that human neuronal networks undergo a structured process of reorganization that leads to the emergence of stable neuronal activity highly resembling the acute state. This study provides a comprehensive characterization of the remarkable adaptive capacity of human neuronal circuits ex vivo. In doing so, it establishes a framework for interpreting functional experiments in human slice cultures and highlights their potential as a translational model system for studying human brain network dynamics.

## Materials and methods

### Patients

The patients were operated at the Clinics of Neurosurgery and Neurointervention, Semmelweis University, 1145 Budapest, Hungary. We received written consent from all patients. Our protocol was approved by the Medical Research Council of the Ministry of Interior (ETT TUKEB, IV-8358/3/2021/EKU and IV-8360/3/2021/EKU) and performed in accordance with the Declaration of Helsinki. Neocortical tissue samples were resected from 30 patients (10 females, 20 males, Supplementary Table 1). We obtained tissue from frontal (*n*=5 patients), parietal (*n*=2) and temporal (*n*=23) lobes. The patients suffered from epilepsy and/or tumour. In case of tumour patients, the resected tissue was always outside of the tumour area.

### Tissue preparation and culturing

Tissue was transported from the operating room to the laboratory and processed for the culturing in an ice-cold, oxygenated NMDG solution^11^. Neocortical slices of 350 µm thickness were cut with a Leica VT1200S vibratome (Leica Biosystems, Buffalo Grove, IL, USA, RRID:SCR_016495). In total, 297 organotypic cultured slices were made from the samples obtained from 31 patients, on average, *n*=10±6 slices were cultured, ranging from 3 to 24 per brain sample. Slice cultures made from one patient were named as HOC (human organotypic culture). Samples obtained from 19 patients were cultured in artificial medium (AM), supplemented with penicillin, streptomycin and amphotericinB (PSA), modified from^11^, slice cultures made from *n*=12 samples were maintained in human cerebrospinal fluid (hCSF), including 7 cultures with only PS, and 5 cultures with PSA supplement (for details see Supplementary material).

### Virus injection

Adeno-associated viruses (AAVs) PHP.eB-EF1a-GCaMP6s-WPRE-pGHpA, (titer: 2.5×10^13^ vg/ml, Addgene #67526), PHP.eB-Syn-jGCaMP7s-WPRE (titer: 2.8×10^14^) vg/ml, Addgene #104487) were injected into *n*=42 slices (derived from 10 patients). At D0, 10 µl was administered into the tissue with a Hamilton-pipette, either at full concentration or diluted to 1:10 or 1:100 in 1xPBS. Slice cultures injected with viruses were distinguished from non-injected slices and were considered as a different HOC sample.

### Recordings

Both acute slices and organotypic cultures were transferred and maintained at 35–37°C in an interface chamber, perfused with a standard physiological solution, and the extracellular local field potential gradient (LFPg) recording was obtained as described previously^12^ (see Supplementary material).

Wide-field imaging was used to assess the calcium (Ca^2+^) signal of GCaMP-positive neurons. Fluorescence imaging was performed on a custom wide-field mesoscope equipped with a tandem lens configuration, utilizing a 50 mm f/1.4 objective lens (Nikon) and a 50 mm f/0.95 imaging lens (Navitar) to yield an effective magnification of 1×. Excitation light from a 470 nm LED (M470L4, Thorlabs) was passed through a 472/30 nm bandpass filter (FF02-472/30-25, Semrock) and coupled into the optical path via a 495 nm dichroic mirror (T495lpxr, Chroma). Emitted fluorescence was collected through a 525/50 nm band-pass emission filter (86-963, Edmund Optics) and detected by a scientific CMOS camera (Prime BSI Express, Teledyne Photometrics). Image acquisition was coordinated using custom Python software interfacing with the camera’s PVCAM library via the PyVCAM wrapper.

### Data analysis

Electrophysiology data were analysed with the Neuroscan Edit4.5 program (Compumedics Neuroscan, Charlotte, NC, USA), and home-written routines for Python 3.14 (Python Programming Language, RRID:SCR_008394).

Detection and analysis of spontaneous population activity (SPA) were performed as described previously^12^ (for details see Supplementary material). Action potentials (AP) of single cells were clustered with a custom-written program for Python (available upon request). Based on their AP width and firing characteristics inhibitory interneurons (IN) and excitatory principal cells (PC) further divided into intrinsically bursting (IB) and regular spiking (RS) types were separated^13^.

Analysis of single cell firing was investigated with the home-written program in Python. For each cell, average firing frequency, inter-spike-interval coefficient of variation (ISI CV), and two measures for burstiness were calculated. When a set of three APs was detected within 20 ms, they were considered to be part of a potential burst. Bursts containing more than three APs could be longer than 20 ms, but each group of three consecutive APs had to lie within a 20 ms period. Burstiness index 1 was determined as the ratio of APs within bursts, whereas burstiness index 2 was the percentage of AP within 20 ms on the autocorrelogram. We also investigated the potential increase in firing rate of each cell compared to SPA events (within ±50 ms of the SPA peak) using an adapted Monte Carlo approach^12^.

Wide-field fluorescence recordings were processed to extract whole-slice and manually defined regional Ca^2+^-transient rates. Following initial quality control to verify temporal consistency, a time-collapsed mean image was generated for each recording, followed by additional image processing steps to improve signal contrast (see Supplementary material). Ca^2+^-transients were automatically identified in the corrected signal using amplitude, prominence, and minimum-distance constraints. Finally, the mean Ca^2+^-transient rate (Hz) was calculated by dividing the total detected peak count by the recording duration.

### Histology

Immunostainings were performed on fixed slice cultures to visualize neuronal and glial presence and morphology, as well as to verify the laminar structure of the neocortex and the cultured slices. Acute and cultured slices used for electrophysiological or wide-field recordings were also fixed after the experiment with a fixative containing 4% paraformaldehyde in 0.1M phosphate buffer (PB), applied for 24 hours. 60 µm thick sections were made from the slices with a Leica VT1200S vibratome. Sections were immunostained against the neuronal cell body marker NeuN, the astroglial marker glial fibrillary acidic protein (GFAP), the microglia marker IBA1, the inhibitory cell markers parvalbumin (PV), calretinin (CR) or the green fluorescent protein for viral expression detection (GFP). For the exact protocol see Supplementary material.

### Quantitative morphological analyses

We determined neuronal density in NeuN-, PV- and CR-stained sections and both astroglial and microglial coverage in GFAP- and IBA1-stained sections, respectively, in slices fixed at different time points, with the aid of the QuPath v0.6.0. software^14,15^. To assess possible statistical differences, we averaged data obtained at D7±1 (week1), D14±1 (week2), D21±1 (week3), D28±1 (week4) and D32 to D45 (week5). For details see Supplementary material.

### Statistics

If the data followed a normal distribution (verified with the Kolmogorov-Smirnov & Shapiro-Wilk tests), a t-test or an unpaired t-test was used to compare two groups. In case of multiple groups, ANOVA (unpaired condition) or ANOVA with Greenhouse-Geisser correction (paired condition) was performed. If the normality test failed, we used Wilcoxon signed rank test (two groups, paired) and Mann-Whitney U test (two groups, unpaired) or Kruskal-Wallis ANOVA (multiple groups, unpaired) and Friedman test (multiple groups, paired) for comparing two or multiple groups, respectively. When comparing multiple groups, Dunn’s post-hoc test with Bonferroni correction was used when the main test reached significance (*p* < 0.05). Significance thresholds were defined as: * *p* < 0.05, ** *p* < 0.01, *** *p* < 0.001. For quantitative histology, we determined statistical significance using GraphPad Prism version 10.0.0 for Windows (GraphPad Software, Boston, MA, USA, RRID:SCR_002798). Statistical analyses of electrophysiological data were performed in Python (v. 3.14) using a custom analysis pipeline, available upon request.

Event-relatedness of each unit was tested via surrogate statistics^13^, which either classified a unit as “increased” (significantly elevated firing during population events) or “indifferent”. Differences in the proportion of event-related units across groups (cell types, days-in-vitro) were assessed using Pearson’s chi-squared test of independence, with results reported as χ², degrees of freedom, and *p*-value.

## Results

### Organotypic slice cultures – culturing conditions

Organotypic cultures were made from human neocortical samples obtained from n=30 patients (Supplementary Table 1, 2). These slices were kept alive up to 42 days in vitro (D42). The viability and activity of the neurons in the cultures were investigated using different methods. In vitro electrophysiological recordings were made to verify neuronal activity both on the day of the operation (D0), as well as at different time points during the culturing period. In case of cultured slices injected with AAVs, neuronal activity was also assessed by wide-field imaging. Furthermore, anatomical techniques were used to estimate the viability and the morphological changes of the slices: neuronal stainings NeuN, parvalbumin (PV) and calretinin (CR) showed the presence and the density of the surviving neurons, whereas glial activation was detected with astrocytic marker GFAP and microglial marker IBA1 stainings. This complex approach helped us to assess the relationship between neocortical activity and the survival of the neuronal and glial networks.

Slices derived from eleven patients were cultured in human cerebrospinal fluid (hCSF), slices obtained from 18 patients were kept in artificial medium (AM), whereas both hCSF and AM were tested in one tissue sample. During the first week of culturing slices were treated with penicillin and streptomycin (PS) or PS completed with amphotericinB (PSA), to prevent bacterial and fungal infections.

### In vitro electrophysiological recordings

Both acute slices and organotypic slice cultures were investigated by in vitro electrophysiological methods. Extracellular recordings were made using a 24-channel linear microelectrode^16^ placed perpendicular to the pial surface to record from the entire width of the cortex. Altogether 90 slices from 17 patients were included in the electrophysiological analysis (D0: 62 slices in 15 patients, D7: 9 slices in 7 patients, D15: 5 slices in 5 patients, D22: 9 slices in 7 patients, D30: 4 slices in 4 patients and D35: 1 slice). Cellular firing as well as the presence of spontaneous population activity^12^ (SPA) emerging in standard physiological baths were noted in all slices (Fig. 1, Supplementary material). In the acute samples (D0) cell firing could be detected in 73% of the slices, 37% of them generated SPA as well. On D7 organotypic slices displayed cellular firing and SPA in 56% and 44%, respectively. Neuronal firing was observed in all D14 slices, of which 60% also produced SPA. On D21, 78% and 33% of slices displayed cellular firing and SPA, respectively. On D30, neuronal firing could be detected in all slices and 50% of them generated SPA as well, and finally the D35 slice showed cell firing without the emergence of SPA. In nine HOC samples we recorded both on D0 and in cultured slices. In five cases SPA was recorded both in acute (D0) and in cultured slices. In three cases SPA could be detected only on D0, but not in the cultures, and in one case SPA was not present on D0 but emerged in all of the cultured slices.

**Figure 1:**
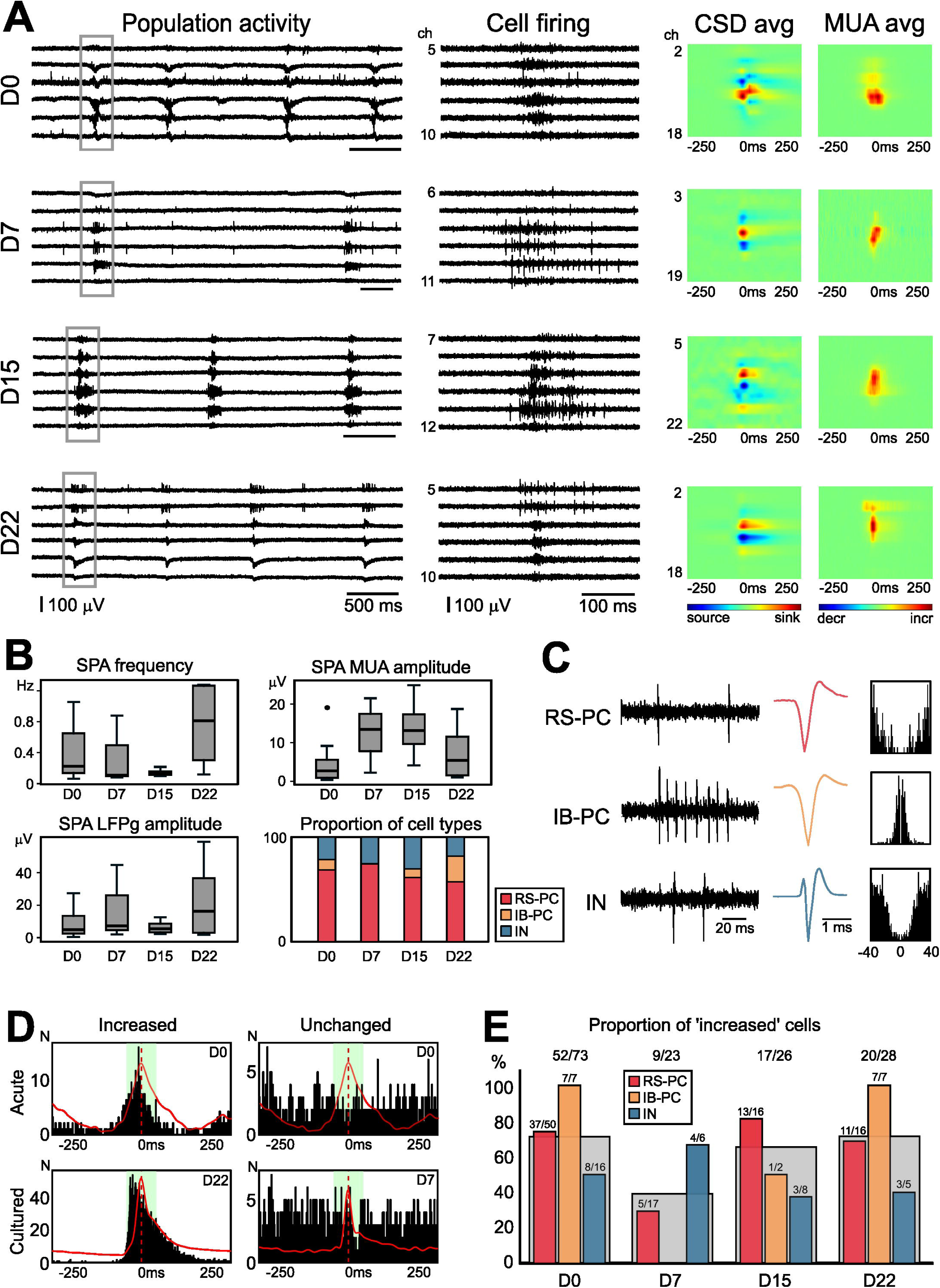
Multiple channel electrophysiological recordings of organotypic cultures. **(A)** Broad-band recordings (left panel) show the spontaneous population activity (SPA) generated in human acute (D0) and cultured (D7, D15, D22) slices. SPA events (grey boxes) consisted of a local field potential transient with increased cell firing. Heat maps generated from current source density (CSD) and multiple unit activity (MUA) analyses show that a complex sink-source pattern and an increased cell firing characterized the events. On the CSD heat maps sinks are marked with warm colours, sources with cold colours. On the MUA maps warm colours represent MUA increase. Heat maps were generated from the time-averaged (avg) SPA. Firing of single neurons (Cell firing) is shown on high pass filtered (500Hz) recordings on the second panel from the left. **(B)** Box plots show the recurrence frequency (left top panel), the LFPg amplitude (left bottom), and the MUA amplitude (right top) of SPAs at the different days of the culturing period. The proportion of the different cell types among the clustered neurons are shown on the right bottom panel. RS-PC: regular spiking principal cell, IB-PC: intrinsically bursting principal cell, IN: inhibitory interneuron. **(C)** High-pass (>500) filtered recordings (left panels), average action potential shape generated from broad band recording (middle panel) and autocorrelogram of one example of RS-PC (top), IB-PC (middle) and IN (bottom) are shown. **(D)** Peri event time histograms demonstrate that clustered single neurons showed either increased or unchanged firing rate during SPA events, both in acute and cultured slices. Red lines show the averaged LFPg of the respective SPA, whereas the green shading marks the duration of the SPA (±50 ms) taken into account in the analysis. **(E)** Proportion of neurons with increased firing rate during the SPAs during the culturing period. Red bars depict RS-PCs, orange bars mark IB-PCs, whereas INs are marked with blue bars. All cells pooled are shown by grey bars.

The recurrence frequency of SPAs in the acute slices (at D0, *n*=11 SPAs from 6 slices from 3 patients, Fig. 1B) was 0.38±0.34 Hz (mean±st.dev.). The recurrence frequency did not change during the culturing period, ranging from 0.05 Hz (D30) to 1.28±0.76 Hz (D22, Kruskal-Wallis ANOVA, *p*>0.05, see Supplementary Table 3). The LFPg amplitude was 94.1±92.1 μV at D0, and was not significantly different during the culturing period (Kruskal-Wallis ANOVA, *p*>0.05), ranging from 41.4 μV at D30 to 233±270 μV at D22. The CSD and the MUA amplitudes were also unchanged on all different days examined (Kruskal-Wallis ANOVA, *p*>0.05). The CSD was 79±67.5 μV at D0 and ranged between 34.6 μV (D30) to 189±213 μV (D22). The MUA was 4.59±5.63 μV at D0 and ranged between 4.12 μV (D30) to 13.9±8.63 μV (D15).

A total of 151 single units (neurons) were clustered in *n*=17 recordings, where SPA was detected (from n=4 patients, Fig. 1C). In the acute slices (4 slices from 3 patients, at D0) a total of 73 neurons were clustered: 57 principal cells (PC, 50 regular spiking, RS, 7 intrinsically bursting, IB) and 16 inhibitory interneurons (IN). In two slices (from 2 patients) recorded at D7, a total of 23 cells were identified: 17 PCs (all RS) and 6 INs. In two slices (from two patients) at D15, we clustered 26 neurons, 18 PCs (16 RS + 2 IB) and 8 INs. In two slices from two patients at D22, 16 RS-PCs, 7 IB-PCs and 5 INs gave a total of 28 cells, whereas we could cluster one IN at D30 (in one slice). The ratio of interneurons did not significantly change during the culturing time (from D0 to D22), it varied between 18 and 31%.

To examine the cellular properties of the clustered neurons (see Methods), we determined the average firing rate, the maximal firing rate, burstiness index 1 (ratio of APs within burst, where burst contains at least 3 APs, is preceded and followed by a 20 ms silent period, and where any three consecutive APs fall within 20 ms), burstiness index 2 (the percentage of ISI values less than 20 ms), half-width 1 (half-width of the largest amplitude of the AP) and half-width 2 (half-width of the second peak, which gives an estimation on the afterhyperpolarization part of the AP), interspike interval coefficient of variation (ISI CV, giving an estimate about the wide or narrow distribution of the ISI values). Higher ISI CV values indicate more irregular firing than lower values.

The average firing rate of all neurons (from D0 to D30) was 1.55±1.92 Hz. The firing rate was similar in the different cell groups (PCs vs. INs, as well as IB-PC vs. RS-PC vs. IN, see Supplementary Material). The average firing rate of all neurons was quite variable along the culturing period: it was significantly higher at D7 and significantly lower at D15 compared to D0 and D22 (see Supplementary Table 4). To examine the effects of culturing on neuronal cellular features, we compared the different cell types clustered in acute slices with those in organotypic cultures (from D7 to D22, pooled). We found changes in several cellular properties affecting different cell types. The firing rate of RS-PCs (but not that of IB-PCs or INs) decreased significantly in culture. The burstiness index 1 of IB-PCs (but not that of RS-PCs and INs) significantly increased in culturing conditions. Note that both burstiness indexes were significantly higher for IB-PCs than the two other cell types, and burstiness index 1 of IB-PCs further increased in culture. Consistent with these groupwise effects, rank-based trend analyses showed modest monotonic tendency across days-in-vitro for IB-PCs (burstiness ISI<20 ms ρ=0.18). Interestingly, the half-width 1 of RS-PCs (but not that of IB-PCs and INs) decreased significantly in culture compared to acute slices. RS-PC spike half-width 1 exhibited a clear negative trend with culture time (ρ=−0.48), consistent with progressive waveform narrowing within this principal cell population. At the same time, half-width 2 did not change for any of the examined cell types. The ISI CV significantly increased for RS-PCs and INs, but not for IB-PCs in culture. When we examined all days, we detected a modest trend in the change of ISI CV (ρ=−0.23, for more details see Supplementary Material).

Next, we verified how the clustered neurons were involved in the population events (SPA) detected in the slice. Therefore, we determined for each cell whether it increased its firing rate during SPA events, with the aid of an adapted Monte-Carlo approach^13^ (see Methods). We could not detect any cells with decreased, only with increased or unchanged firing rate (Fig. 1B, D, E). In the acute slices 53/73 cells (71%) increased their firing rate, whereas the others remained unchanged. The ratio of participating cells (with increased firing rate) was significantly different during the culturing period: 9/23 (39%) cells at D7, 17/26 (65%) neurons at D15 and 20/28 (71%) cells at D22 increased their firing rate during SPA events (Chi-square test, *p*=0.009). When regarding the involvement of the different cell types, we found that all three neuron groups participated in the generation of SPAs, although at a significantly different ratio (67% of RS-PCs, 94% of IB-PC and 50% of INs, Chi-square test, *p*=0.036, Fig. 1). Within-cell-type comparisons indicated that responsive (“increased”) RS-PCs had higher ISI<20 ms burstiness than non-responsive RS-PCs (*p*<0.001), whereas responsive interneurons fired faster (*p*=0.022) and were less bursty (*p*=0.040), than INs with unchanged firing rate.

### Wide-field imaging

To assess neuronal activity in the organotypic slice cultures, we performed wide-field imaging of slices expressing GCaMP following viral transduction (Methods, Supplementary material). Seventeen slices (from three patients) were recorded across D1 to D31 post-injection. Neuronal GCaMP expression was confirmed in 15 of 15 injected slices, whereas two non-injected slices served as negative control and showed no fluorescent transients.

GCaMP signal was first detected on D4 in slices receiving undiluted virus, and remained recordable until D31. Across the 14 slices with longitudinal recordings fluorescence intensity and spatial extent increased over time in the majority of the cases (10/14 slices, Fig. 2, Supplementary Fig. 2A, D, G).

**Figure 2:**
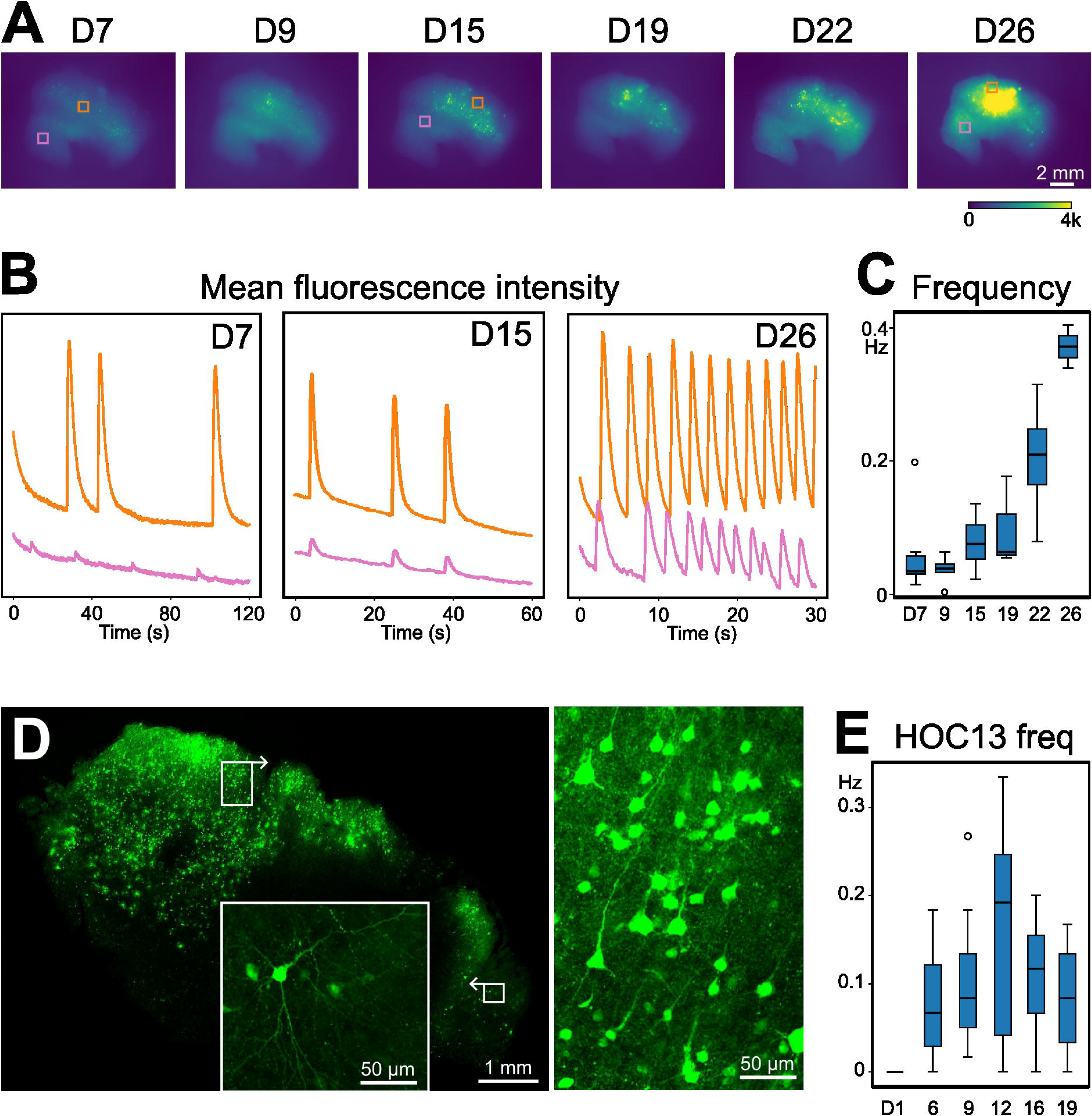
Wide-field Ca^2+^-imaging reveals spatially confined SPA over time in virally transduced human organotypic slices. **(A)** Time-averaged fluorescence intensity images of a single organotypic slice (HOC15_p1w3, undiluted virus) at different time points of the culturing period (from D7 to D26). The slice was injected with EF1a-GCaMP6s-WPRE (PHP.eB, titer: 2.47E+13 vg/ml) and fixed on D28. Warmer colours (green-yellow) show higher fluorescence (arbitrary unit, 0-4000), same colour scale applies to all images. **(B)** Fluorescence intensity traces (arbitrary units) from two spatially distinct region of interest (orange and pink squares in A) at D7, D15 and D26. Peaks correspond to population Ca^2+^-transients. SPA events are not consistently synchronous between regions, indicating multiple hubs of activity generation. Note that time axes differ between recording days and are indicated below each trace. **(C)** Ca^2+^-transient recurrence frequency measured over the full field of view on the slice in (A) accross the culturing period. Consistent with the traces shown on (B), the frequency gradually increases with time spent in culture. **(D)** Confocal image of the slice shown on (A) and (B) demonstrate the presence of numerous GCaMP-positive cells. Numerous cells are present all over the whole slice, although with variable density. Boxed regions (upper, lower) are shown at higher magnification in the right panel and inset, respectively. **(E)** Ca^2+^-transient recurrence frequency of SPA across the culturing period for all slices (n=6) from one HOC sample. Note that prior to GCaMP expression, at D1, zero frequency reflects reporter absence. Consistent detection of transients across slices from D6 onward indicates reliable reporter expression and recurrent SPA with stable frequency over the culture period.

Population Ca^2+^ activity – the correlate of SPA recorded with electrophysiology – was quantified by GCaMP-reported transients over the full slice. All 15 GCaMP-positive slices showed circumscribed regions of synchronous Ca^2+^ fluctuations on at least one recording day, and negative controls showed none. These active spots usually remained spatially stable during the culturing time and could be observed at every imaging occasion, up to D31. GCaMP-related activity appeared as first at D4 (n=2/6 slices) and was sustained in 11 of 15 slices (73%) for at least 19 days post-injection. The temporal profiles of Ca²C-transient frequency varied across slices and across patients. The frequency gradually increased with time in most slices (n=5, Fig. 2B, Supplementary Fig. 2A), only one slice showed a decreasing pattern. In two slices the frequency was low at the early period (up to D9), then stayed high until the end of the culturing time (see Supplementary Fig. 2E). Further three slices showed a biphasic pattern: increasing frequency to D9-D20, which declined at the end (D22-D31) of the culturing period (see Supplementary Fig. 2B). In three slices from the same patient the recurrence frequency of the population events was stable all over the entire culturing time (see Supplementary Fig. 2H).

### Quantitative morphological analysis

To characterize the morphological changes in the human neocortical organotypic cultures we performed quantitative histological analysis on 280 slices derived from 28 patients (Supplementary Table 2). using NeuN staining to assess neuronal density, GFAP and IBA1 to estimate astroglial and microglial activation, respectively, as well as parvalbumin (PV) and calretinin (CR) immunolabeling to quantify inhibitory interneuron subtypes. Culturing produced consistent and substantial structural remodeling: neuronal density considerably declined, both inhibitory interneuron subtypes were significantly reduced, and reactive gliosis was prominent throughout the culture period. Unless otherwise stated, data from epileptic and tumour-associated samples were pooled where no significant group difference was found. Effects of culturing conditions — medium type (human cerebrospinal fluid, hCSF vs. artifical medium, AM), antimicrobial supplement (penicillin+streptomycin, PS vs. PS+amphotericinB, PSA), and viral treatment — are reported separately where applicable. One human organotypic culture (HOC) sample was considered as the sum of the slices made from a tissue sample derived from one patient, including both acute slices (D0) and cultured slices (from D1 up to D42). We hypothesized that virus injection affects the morphological status of the organotypic slices. Therefore, slices injected with viruses were treated separately, and were considered as separate HOC samples, even if they derived from the same patient.

### Neuronal loss in human organotypic cultures

The acute neocortical samples (at D0) showed the typical 6-layered structure of the human neocortex in all cases. Compared to D0, the laminar architecture of the neocortex has been notably changed during culturing (Fig. 3). The neuronal density visibly decreased over time, although the most pronounced cell loss occurred during the first week. While all six neocortical layers were visible in the acute tissue, after D7 the layer borders started to vanish. Intracellular modifications were also observed with time. Compared to the more homogenous staining, NeuN was mainly localized within the nucleus and the cytoplasm became lightly labelled (Fig 3). Neuronal debris appeared in the neuropil already after the first week. These changes occurred earlier when slices were cultured in hCSF. The use of AM resulted in a better preservation of the neurons and the cortical structure, decrease of the neuronal cell numbers and the appearance of the debris (Supplementary Fig. 3). The neuronal survival was variable during the time course of the culturing period: large differences were seen in the neuronal density, in the integrity of the cells (lack of NeuN-positive debris), which could affect either supragranular or infragranular or both layers. The background of the sections became considerably brighter with time, however this change was also very variable among samples.

**Figure 3:**
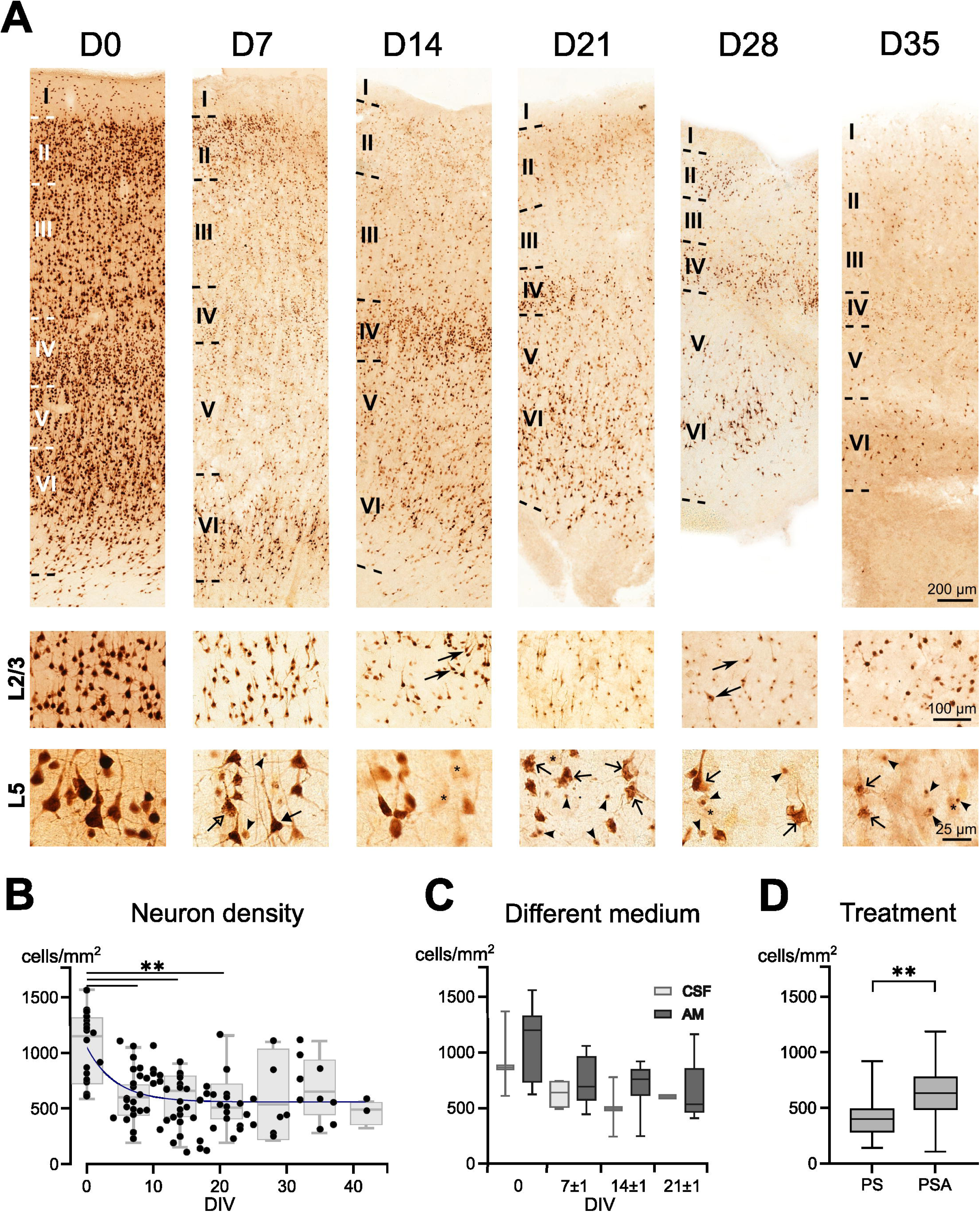
Considerable neuronal death occurs in organotypic cultures. **(A)** The cortical structure of human organotypic slices is preserved during the culturing period. The upper row of pictures shows the entire cortex including all visible layers, the middle row shows the neuron density with higher magnification, whereas on the bottom row the morphology of the neurons can be seen. The human neocortex shows the typical 6-layered structure in acute slices (D0) in NeuN-stained sections. At D7, all 6 layers can be distinguished, but after the second week (from D14), layer borders usually vanish due to neuron loss. At D0 neocortical neurons show normal morphology: the nucleus is dark, and NeuN is visible in the proximal dendrites (middle and bottom). After one week of culture, the background of the NeuN-stained section starts to clear up, but most neurons still show normal morphology (arrow on bottom image). However, cells with inhomogeneous NeuN-staining (triangle arrow) and NeuN-positive cellular fragments (open arrow) appear. In several cases, the surviving neurons were organized in columns (left side of D7 bottom image). After the considerable cell loss of the first week, the orientation of the surviving neurons was often changed, the apical dendrite of the pyramidal cells arose towards different directions (sharp-edged arrows on D14 middle image). Furthermore, numerous neurons were only lightly stained (asterisks on bottom images), and more neuronal fragments, debris (arrowheads on bottom images), and neurons with distorted cell body and dendrites (open arrows on bottom images) could be detected. **(B)** Neuronal density shows a significant decrease during the first two weeks of culture, then it stays stable. Black dots represent the neuron density of each examined slice, fixed at different time points, whereas grey boxplots show data averaged at the end of each week (±1 day). Neuron density at the end of the first, second and third weeks was significantly different from D0 (** *p*=0.0006). **(C)** No significant differences were found when comparing neuron densities in slices kept either in human cerebrospinal fluid (CSF) or in artificial medium (AM). However, note that cell densities are higher in AM. **(D)** Antibiotic treatment (Penicillin+Streptavidin, PS) resulted in significantly lower neuron densities than antibiotic+antifungal (PS+amphotericinB, PSA) treatment.

We performed cell counting on NeuN-positive sections of *n*=118 organotypic slices, ranging from D4 to D42, plus *n*=17 D0 slices, derived from a total of 25 patients (for exact numbers of slices see Supplementary Table 2, for cell density values see Table 1 and Supplementary Table 9). There were no significant differences between the examined 34 HOC samples (28 control samples, 6 virally injected samples, Kruskal-Wallis ANOVA, *p*=0.0805). No differences were seen when comparing epileptic to non-epileptic samples at D0 (unpaired t-test, *p*=0.1506) and thus, the data derived from epileptic and tumour patients was pooled for further analyses. The cell density significantly decreased over time in samples without viral injections (day-by-day changes, Supplementary material, Kruskal-Wallis ANOVA, *p*=0.0105, Fig. 3B). Significantly reduced cell density was observed when examined at the end of every week (week-by-week analysis, see Supplementary Table 9), i.e. at D7±1, D14±1 and D21±1 compared to D0 (ANOVA, *p*=0.0006).

**Table 1.** Quantitative histological data. Neuron densities (cells/mm^2^) are shown for NeuN, PV and CR stainings. Glial coverage (%) are given for GFAP and IBA1 stainings. Furthermore, the perimeter-to-area ratio (PAR) is also given for IBA1+ cells. All data are given in median [1^st^ quartile 3^rd^ quartile] and mean±standard deviation

|  |  | D0 |  | D7±2 |  | D14±2 |  | D21±2 |  | D28±2 |  | D33+ |  |
| --- | --- | --- | --- | --- | --- | --- | --- | --- | --- | --- | --- | --- | --- |
|  |  | n | value | n | value | n | value | n | value | n | value | n | value |
| NeuN density (cells/mm<sup>2</sup>) | median [Q1 Q3] | 17 | 1116 [761 1291] | 22 | 602 [488 789] | 28 | 567 [414 815] | 19 | 536 [392 690] | 9 | 442 [327 804] | 13 | 585 [357 697] |
|  | mean±st.dev |  | 1068±333 |  | 640±234 |  | 598±248 |  | 599±320 |  | 590±341 |  | 621±399 |
| GFAP coverage (%) | median [Q1 Q3] | 20 | 36.98 [25.85 52.65] | 19 | 19.53 [10.81 33.43] | 29 | 30.41 [22.52 44.92] | 13 | 41.47 [19.52 54.20] | 7 | 27.24 [17.30 29.25] | 12 | 36.62 [21.37 43.33] |
|  | mean±st.dev |  | 39.35±17.84 |  | 23.34±16.51 |  | 32.78±15.95 |  | 36.47±20.60 |  | 26.88±13.28 |  | 34.87±16.98 |
| IBA1 coverage (%) | median [Q1 Q3] | 11 | 19.22 [13.78 38.39] | 5 | 17.15 [11.14 17.65] | 9 | 11.47 [9.18 28.26] | 7 | 25.55 [13.52 27.68] | 1 | 21.04 | 9 | 34.04 [26.27 40.74] |
|  | mean±st.dev |  | 29.72±23.50 |  | 16.12±7.78 |  | 16.44±9.76 |  | 21.38±8,64 |  | 21.04 |  | 37.99±18.95 |
| IBA1 PAR | median [Q1 Q3] | 11 | 1.94 [1.82 2.37] | 5 | 4.07 [3.59 4.10] | 9 | 4.03 [3.89 4.55] | 7 | 3.93 [3.70 4.06] | 1 | 4.04 | 9 | 3.52 [3.09 3.62] |
|  | mean±st.dev |  | 2.09±0.37 |  | 3.86±0.80 |  | 3.96±0.89 |  | 3.90±0.22 |  | 4.04 |  | 3.43±0.36 |
| PV density (cells/mm<sup>2</sup>) | median [Q1 Q3] | 14 | 25 [19 34] | 9 | 2 [1 20] | 11 | 2 [0 6] | 7 | 6 [0 8] | 4 | 14 [9 18] | 14 | 2 [0 6] |
|  | mean±st.dev |  | 32±24 |  | 12±14 |  | 6±10 |  | 7±9 |  | 13±7 |  | 4±6 |
| CR density (cells/mm<sup>2</sup>) | median [Q1 Q3] | 11 | 69 [39 92] | 7 | 27 [5 48] | 10 | 18 [10 35] | 8 | 12 [4 23] | 4 | 9 [7 13] | 8 | 8 [2 37] |
|  | mean±st.dev |  | 70±46 |  | 34±38 |  | 26±22 |  | 14±11 |  | 11±9 |  | 18±20 |

No statistical difference was observed, when we examined slices with or without viral infection derived from the same patients (Wilcoxon test, *p*=0.9344). Despite the considerable differences in neuronal morphology and in the cortical laminarization changes in slices kept in hCSF or AM (see Supplementary Fig. 3, Supplementary Table 9), we could not show significant differences in the neuronal density (Fig. 3C, unpaired t-test, *p*=0.1285). The use of antibacterial PS treatment resulted in significantly lower values in the neuronal density, than that in slices with complex antibacterial+antifungous PSA treatment (Fig. 3D, unpaired t-test, *p*=0.0075).

### Astroglial activation

To examine the changes in the astroglial network, and to reveal the formation of a possible glial scar, we performed glial fibrillar acidic protein (GFAP) staining on *n*=104 organotypic slices plus 20 samples fixed at D0, derived from 25 patients (Supplementary Table 2). In about half (11 of 20) of the acute slices (D0), the neocortex GFAP+ cells showed the features of resting astroglial cells. In the remaining samples gliosis was observed and activated astroglial cells were visible either in patches or extending to the entire slice. Gliosis was not related to epilepsy, as it occurred in samples derived from both epileptic (*n*=4) and non-epileptic tumour (*n*=5) patients. Six out of 20 slices were also used for electrophysiological recording, and four of them were gliotic. During the culturing period, we detected a dramatic change in the astroglial morphology. Astroglial cells became very heavily stained, the size of their cell body increased, and their processes became thick (Fig. 4). Blood vessel endfeet were usually strongly stained, although the vessels were most likely not functional in organotypic slices (see Supplementary Fig. 4). Different astroglial patterns were observed in the cultures. Either scattered glial cells were visible with a very bright, cleaned background, or a very dense glial meshwork covered part or the entire area of the slice. Reorganisation of GFAP+ network within the three-dimensional structure of the organotypic slice showed large differences: a glial scar formed on the top and the bottom of the slice, whereas in the middle only moderate gliosis was observed (Fig. 4). In several cases we saw long glial processes stretching from layer 1 into the deeper layers, where a scarce network of glial cells was visible. This phenomenon was most prominent in virally injected slices, but was also visible in non-injected cultures as well (Supplementary Fig. 4F).

**Figure 4:**
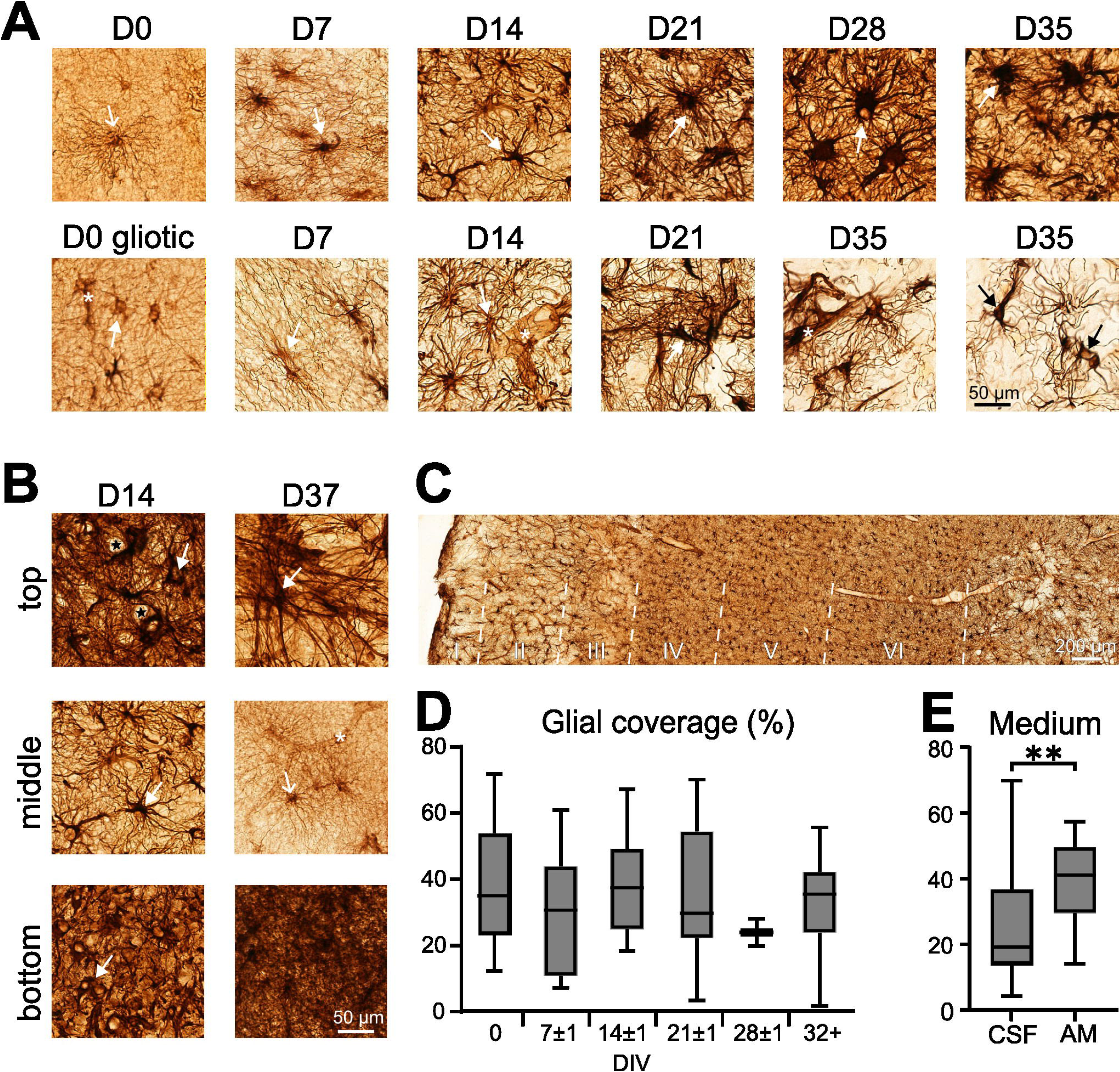
GFAP-positive astroglial cells are highly activated in organotypic cultures. **(A)** Morphology of GFAP+ astroglial cells during the course of the culture period. In the acute tissue, GFAP+ cells showed either the morphology of resting astroglial cells (D0, open arrow), or the signs of gliosis (D0 gliotic), where GFAP+ activated astroglial cells possessed larger cell body and thickened processes (arrow), both types forming the typical endfeet around the blood vessels (asterisks). During the culture period, activated astroglial cells were visible in all cases, resting astrocytes were very rarely seen. GFAP staining was quite variable, it became either very dark with large, dark cell bodies and numerous thick and heavily stained processes (arrows on upper row of images), or the background became clear and GFAP+ activated astroglial cells showed a less dense distribution (bottom row of images), and numerous thin processes. Blood vessels (asterisks) were usually drawn by the astroglial endfeet, even at D35. **(B)** Heavily stained glial scar was formed on the two surfaces of the slices (top and bottom), whereas moderate (D14) or little (D37) gliosis was observed in the middle of the slices. Arrows point on activated astroglial cells, an open arrow shows a resting glial cell (D37, middle), whereas an asterisk marks a long blood vessel present at D37. Empty holes (black stars, D14 top) or long glial processes (D37 top) were present in some cases on the top the slices. Note that the GFAP staining is the strongest on the bottom of the slices. **(C)** GFAP-staining delineates the layers of the neocortex in a D14 organotypic culture. A very dense and dark network covers the granular (L4) and infragranular layers (L5-6), whereas activated astrocytes form a looser network in the supragranular layers (L1-3) and in the white matter. **(D)** Glial coverage did not significantly change during the culturing period. Box plots show averaged data at the end of each week (±1 day). Note that the glial coverage showed a high variability among the samples all over the culturing period. **(E)** Significantly lower glial coverage was seen in slices kept in human CSF compared to AM.

Gliosis was quantified by determining glial coverage within the slices (Methods, Table 1, Supplementary Table 9). In general, glial coverage showed a high variability between sections. Glial coverage was not significantly different between the HOC samples (Kruskal-Wallis ANOVA, *p*=0.1316), neither between slices treated either with PS or with PSA (unpaired t-test, *p*=0.9718). Despite the striking modifications in the morphology of the astroglial cells, glial coverage was not different during the culturing period (day-by-day changes, Kruskal-Wallis test, *p*=0.1644), neither when we compared the end of each week to D0 (ANOVA, *p*=0.5611). Slices kept in hCSF showed lower glial coverage than those cultured in AM (unpaired t-test, *p*=0.0045), but viral injection had no effect on the glial coverage (Wilcoxon test, *p*=0.3125).

### Microglial activation and morphological heterogeneity

The morphology of microglial cells is linked to their functions. Ramified appearance is characteristic to homeostatic resting state when they surveille the neighbouring neurons. Rod-like and honeycomb morphologies suggest injury-induced activation, hyper-ramified forms appear in stress, whereas reactive, jellyfish and ball-like amoeboid types are known to show a phagocytic function^17–19^. To assess the changes in the microglial network in organotypic cultures, IBA1 staining was performed on a total of *n*=40 organotypic slices plus 10 samples fixed at D0, derived from a total of 13 patients (Supplementary Table 2). In almost half of the acute samples (D0), IBA1-positive microglial cells were distributed homogeneously and showed the homeostatic ramified morphology (*n*=4/10 samples). In the remaining six cases, activated microglial morphologies were seen, such as hyper-ramified (*n*=4/6 samples), reactive (*n*=2/6) and elongated amoeboid-like (*n*=1/5) microglial cells. Rod-like microglia were visible together with homeostatic, ramified cells in two cases, whereas they were present with other activated cell types in six cases. All sections containing homeostatic ramified microglia cells derived from therapy resistant epileptic patients, whereas the samples with activated microglial cells derived mainly from patients with tumour (5 tumour patients, 1 epileptic patient). In the cultured slices drastic morphological changes occurred. The distribution of the microglial cells was inhomogeneous, both spatially (Fig. 5A), and morphologically (Fig. 5B). The large differences in the density of the IBA1+ cells - such as in the case of GFAP+ astroglial cells - was also linked to the top-bottom gradient of the slice (Supplementary Fig. 5), i.e. dense networks of activated microglia were present at the surface of the slice, whereas the density of microglial cells was less in the middle of the slice. The appearance of the IBA1+ cells was extremely variable within and among the slices and patients. The size of the cell body ranged from small to giant, the number of processes spread from zero to very high numbers. Furthermore, the thickness and the length of their processes were also considerably variable, such as the intensity of their IBA1-staining, in all different combinations of all the above features. In culture, homeostatic resting types totally disappeared, while all other types were detected. The ratio of the slices showing rod-like, honeycomb (related to injury), hyper-ramified (linked to stress), and ball-like amoeboid, jellyfish and reactive-hypertrophic types (related to phagocytosis) were determined during the course of the culturing period (Fig. 5E). Rod-like and hyper-ramified forms were usually seen at the earlier phase of the culture period (D7-D21), but the appearance of these types could be associated with viral injection at later stages (D32-D37) as well. Honeycomb microglia appeared mainly at later phases, both in virally injected and non-injected slices (D32-D42). Phagocytic reactive, jellyfish and amoeboid were the most frequently seen forms, through the entire culturing period. In several cases, different forms of microglial cells, such as amoeboid and reactive types formed large groups, and were located next to each other (Supplementary Fig. 5A-C). In organotypic slices kept in hCSF amoeboid/phagocytic forms were more frequently seen, whereas in AM, more microglial cells showed stress related activation, i.e, had smaller cell body and had numerous long, thin processes (hyper-ramified type).

**Figure 5:**
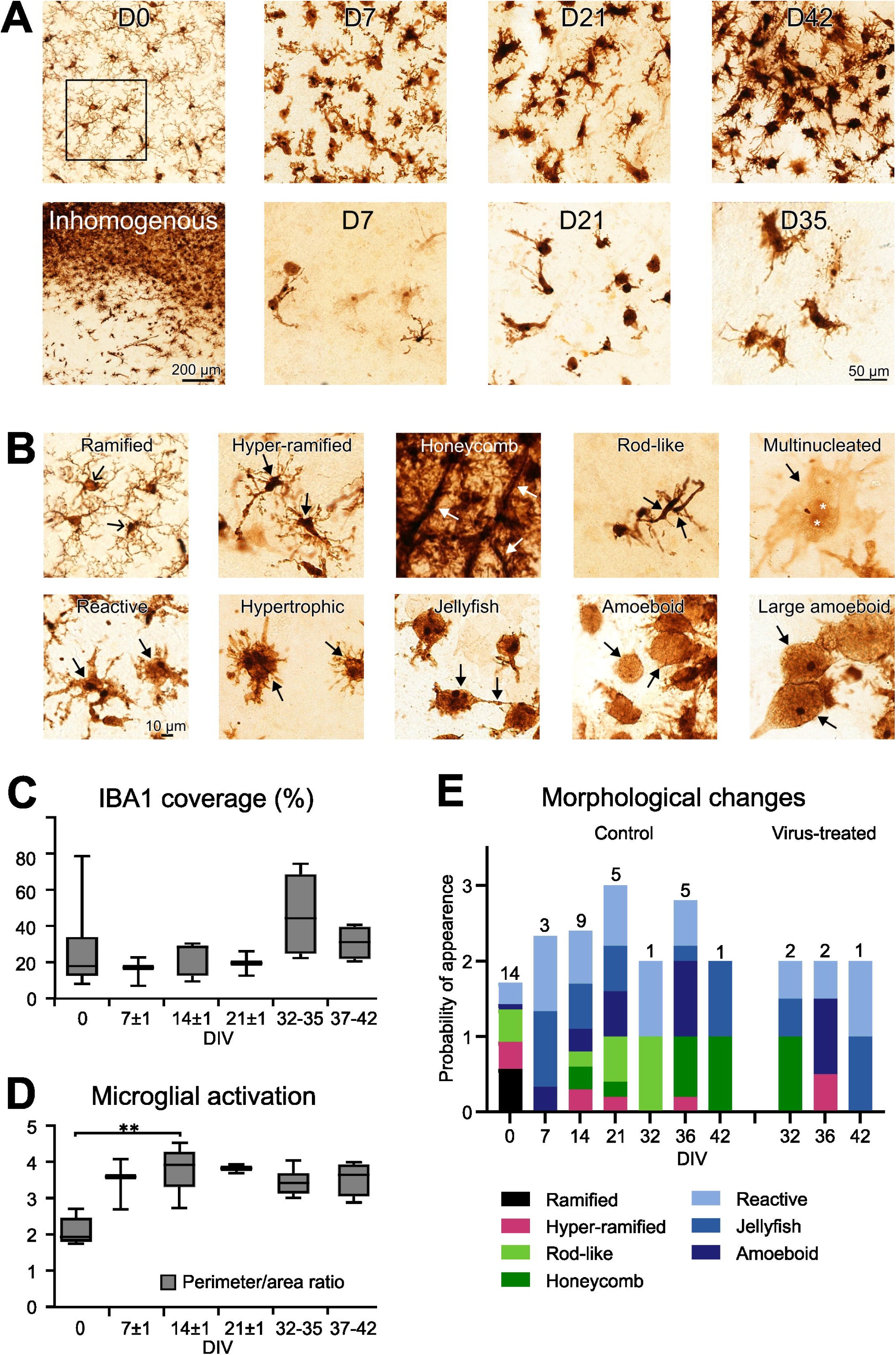
IBA1-positive microglial cells show a dramatic change in human neocortical organotypic cultures. **(A)** Morphology and distribution of IBA1+ microglial cells. At D0, resting microglial cells were visible. After D0, only activated microglial cells were observed, resting IBA1+ cells disappeared. Areas with high (upper row) and low (lower row) density of microglial cells were both present during the entire culturing time (D7 to D42). Patches of high and low density IBA1-stained areas could be detected next to each other, in the same slice (Inhomogeneous). **(B)** Microglial cells showed very variable morphology. Resting cells have ramified morphology (open arrows), i.e. small cell body and several ramifying processes. Black arrows show all different types of activated glial cells on the images. Hyperramified cells have small cell bodies with numerous thin processes. Honeycomb structures are composed of several cells with large and dark processes (white arrows). Rod-like cells have small and elongated cell bodies and only a few (if any) processes. We have seen multinucleated cells with giant cell bodies (white asterisks show the nuclei). Usually, these cells were lightly stained. Reactive microglial cells have large cell bodies with several thick processes. Hypertrophic cells possess very large cell bodies and numerous short processes, whereas jellyfish type microglia usually have only few short processes. Amoeboid and large amoeboid cells have round giant cell bodies without processes. Note that the same scale bar applies to all images. **(C)** Microglial coverage of the organotypic slices did not change significantly during the culturing period. Note the large variability of the IBA1 coverage at D0. DIV: days in vitro. **(D)** The perimeter/area ratio showed that a significant microglial activation occurs during the first 2 weeks. The perimeter/area doubled compared to D0 and remained high during the culture period. ** *p*<0.01 **(E)** Probability of appearance of the different types of microglial cells during the culture period. The ratio of slices containing the different microglia types was determined at each examined day (DIV) and added. The numbers of examined slices are shown above the bars. Virus-injected slices (right) were examined separately. Ramified microglial cells (black) were found only at D0, in about half of the slices. Injury-linked hyper-ramified types (magenta) were seen relatively rarely. Stress-related honeycomb and rod-like microglia (green bars) were seen from the second week. The phagocytic types (amoeboid, jellyfish and reactive, blue bars) were present during the entire culture period.

To quantify changes in microglial activation during the culture period, we determined IBA1 coverage (percentage of the examined slice area) and calculated the perimeter-to-area ratio (PAR) of IBA1+ cells within predefined areas (see Methods, Table 1, Supplementary Table 9). Despite the considerable morphological changes over the culturing period, microglial coverage did not show significant modifications (Kruskal-Wallis ANOVA, *p*=0.3015). Furthermore, we could not demonstrate significant differences between the tissue samples derived from different patients (*n*=13 samples without viral treatment, *n*=37 slices, Kruskal-Wallis test, *p*=0.3566), nor when comparing different culture conditions (hCSF vs AM: Mann-Whitney test, *p*=0.6620; viral treatment vs. not: paired t-test, *p*=0.9296). However, the difference in microglial coverage between the middle and the surface of the slices increased to 6-fold over 6 weeks of culturing, as also shown by the increase in standard deviation (see Supplementary material).

In accordance with the coverage, the PAR did not show significant differences between the cultures (Kruskal-Wallis test, *p*=0.2438) nor over the culturing period (Kruskal-Wallis test, *p*=0.5553) nor when slices were treated with viruses (Wilcoxon signed rank test, *p*=0.25). However, the week-by-week changes were significantly different (Kruskal-Wallis ANOVA, *p*=0.0029). After the second week, the activation level of microglial cells - shown by the PAR - significantly differed from the original state (multiple comparison Dunn’s test, *p*=0.0047). In AM, the PAR was significantly lower compared to hCSF (Mann-Whitney test, *p*=0.0027).

### Loss of parvalbumin-positive perisomatic inhibitory cells

Parvalbumin-stained neurons are known to be perisomatic inhibitory interneurons in the human neocortex^20^, comprising basket and axo-axonic cells^21,22^. Light microscopic analysis was done in 59 organotypic and 14 acute (D0) slices stained against PV, derived from 20 patients. In most of the acute postoperative tissue samples numerous PV+ cells were visible in neocortical layers 2-6, with a dense axonal network in layer 4 (Fig. 6A). The multipolar cells usually possessed long, well-stained dendrites, oriented mainly perpendicular to the pial surface. The number of PV+ interneurons was reduced in organotypic cultures, already at D7, and remained very low throughout the culture period. Usually, in samples derived from epileptic patients PV+ cells remained visible for a longer period than in tissues from tumour patients. PV+ cells in L4 were preserved for the longest culture period and remained visible even at D35-D42. At the beginning of the culture period (first two-three weeks), the majority of the surviving PV+ cells showed similar morphology to the acute tissue (D0), but cells with large cell bodies and numerous long dendrites also appeared (Fig. 6B). From two-three weeks in culture, we observed regions in the slices which lacked PV+ cell bodies, together with preserved axonal clouds (Fig. 6B, C). At the end of the culture period (fourth to seventh week) the number of PV+ cells dropped close to zero, and large areas were void of stained elements. In these cases the neocortical layers were not distinguishable (Fig. 6C). The remaining cells were usually located in L4 and possessed large cell bodies and distorted dendrites. The axon terminal structures typical to axo-axonic cells (the so-called “chandeliers”), as well as the typical basket-like formations characteristic to basket cells were visible all over the culture period (Fig. 6A1–C2).

**Figure 6:**
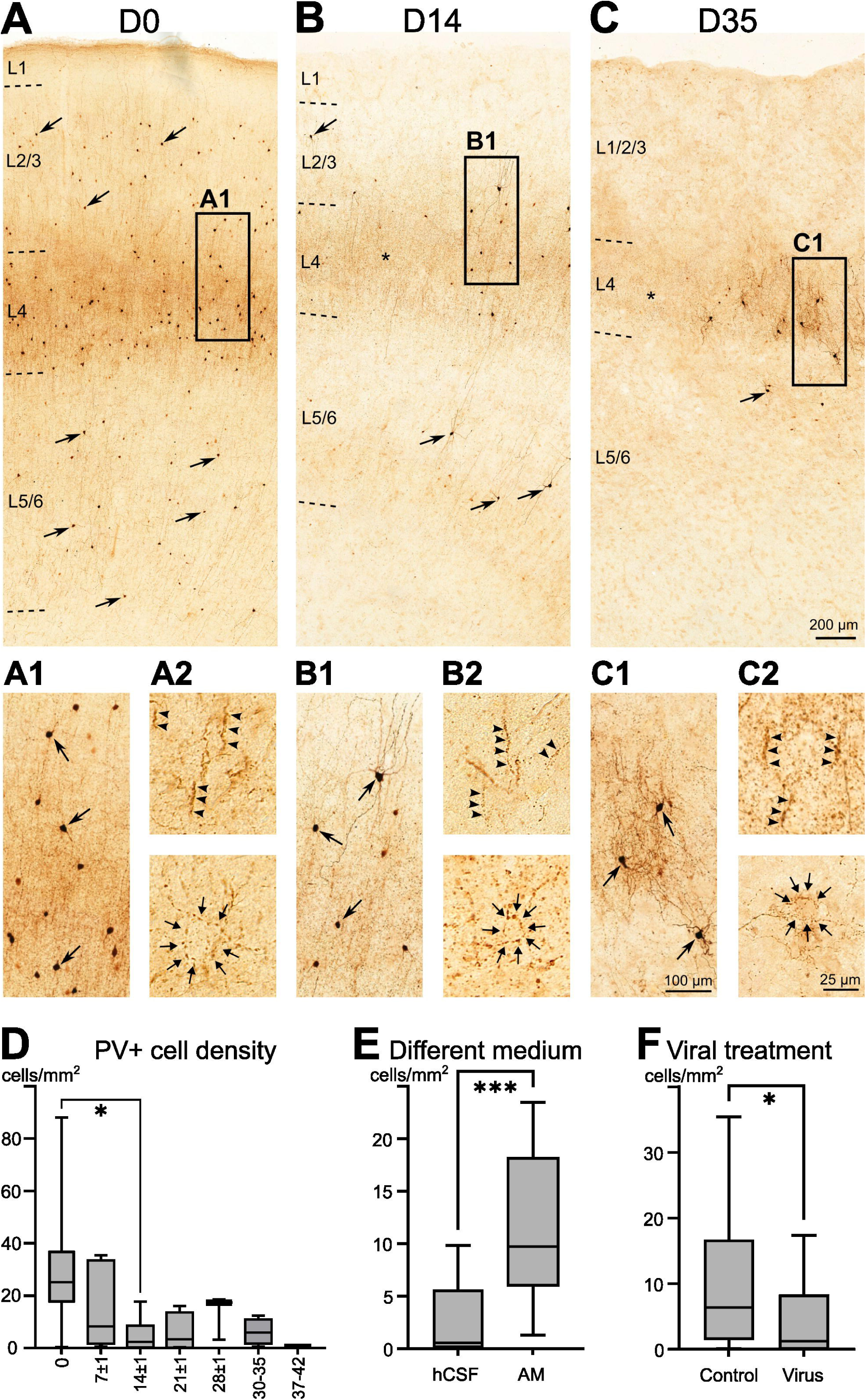
PV-stained cells are mainly lost in organotypic cultures, but the remaining ones preserve their functions. **(A)** Parvalbumin stained section from an acute sample. Numerous PV+ interneurons (arrows) are located in all neocortical layers except L1. A dark axonal band delineates L4. The area in the black box is magnified in A1. Note the dense PV+ axonal staining and the large number of cell bodies. A2) The typical axo-axonic (top, arrowheads) and basket (bottom, small arrows) axonal structures are visible in the acute samples. Axo-axonic formations were the most prominently visible in L5/6. **(B)** The number of PV+ neurons has significantly decreased in D14 cultures, in all layers. The laminar structure of the neocortex is still visible, the layers can be distinguished. Note that only scattered PV+ interneurons are visible in L2/3 and L5/6. In L4, the loss of PV+ cells occurred in patches, the boxed area is magnified on B1. The neighbouring area marked with an asterisk is lacking PV+ cells, while the axonal band is still visible. **(B1)** At D14 several cells with a considerably large cell body and long and numerous dendrites appear (upper cell marked with an arrow). **(B2)** The typical axo-axonic (top, arrowheads) and basket (bottom, small arrows) formations are present in L5/6. **(C)** The number of PV+ interneurons has dramatically decreased at the end of the culturing period. In this sample from a D35 culture, cell bodies were present only in patches, mainly in L4. The axonal band still delineates L4, but the borders of other layers cannot be distinguished. The axonal band is present even in regions without PV+ cell bodies (asterisk), although less darkly stained than at earlier stages of culture. **(C1)** The box on the upper image is magnified. The remaining cells usually have a large cell body and numerous tortuous dendrites (arrows). **(C2)** The axo-axonic (top, arrowheads) and basket formations (bottom, small arrows) are still present. **(D)** PV+ cell density was reduced during the culture period. The numbers of PV+ neurons dropped at the beginning, and then stayed low during the later period of culturing. The PV+ neuron density was significantly lower at D14 than at D0. * *p*=0.0283 **(E)** The use of the different cell culture medium had a significant effect on the PV+ cell density. It was significantly higher in artificial medium (AM) than in human cerebrospinal fluid (hCSF). *** *p*=0.0005 **(F)** Viral treatment has a significant effect on the PV+ cell density, which was significantly lower in virus treated samples compared to non-treated control. * *p*=0.0448

The change in PV+ cell density was quantified in acute and cultured slices (see Methods, Table 1, Supplementary Table 9). The origin of the tissue did not impact the cell density in samples without viral injections (Kruskal-Wallis ANOVA, *p*=0.5206), and there was no significant difference between epileptic and non-epileptic tissues at D0 (unpaired t-test, *p*=0.4912). Among the different culturing conditions, the medium type and the viral treatment caused significant differences (hCSF vs. AM: Mann-Whitney test, *p*=0.0005; viral treatment vs. control: Mann-Whitney test, *p*=0.0448); the presence of the antimycotic agent did not affect the cell density during the culturing in CSF (PS vs. PSA, Mann-Whitney test, *p*=0.0697). Considering the change in cell density across all measured samples over the whole culturing period (D0 to D42), the difference was significant (Kruskal-Wallis ANOVA, *p*=0.0194), such as in the case of the week-by-week comparison (Kruskal-Wallis ANOVA, *n*=44 slices, *p*=0.0082). The most remarkable difference was between the D0 and D14 state (post-hoc test: *p*=0.0283). When examining the day-by-day changes only in cultures without viral injection (from D0 to D42), we found no significant differences in the PV+ cell density (Kruskal-Wallis test, *p*=0.1044).

### Loss of calretinin-stained neurons

Calretinin-positive inhibitory neurons in the human neocortex are known to be either inhibitory interneurons found in each layers of the neocortex^23^, or Cajal-Retzius cells located in layer 1^24^. To assess modifications in CR+ cells, we analysed CR-stained sections from 43 organotypic slices and 11 acute samples spanning D0-D42 obtained from 11 patients. In the acute samples (at D0) numerous immunostained cells were visible in all layers of the neocortex. In layer 1, large bipolar horizontal cells were present, which were identified as Cajal-Retzius cells (Fig. 7A, B). In the other layers (L2-L6) bipolar and multipolar cells with considerably smaller cell bodies were observed. These cells were usually perpendicularly oriented to the pial surface, and frequently possessed parallelly running dendritic bundles (Fig. 7D). Calretinin-positive cells in epileptic tissue usually had a more complex dendritic tree than cells in tumour tissue, and they preserved this feature throughout the culturing period. Beaded dendrites were observed in several cases even in the acute samples. During the culturing period, the number of CR+ cells has gradually decreased. However, the decrease was not homogeneous, after the second week of culture slices with high and low numbers of CR+ interneurons were seen, sometimes derived from the same HOC sample, or patches of high cell density and empty areas in the same slice (Fig. 7A). At the late stages of the culturing period (after D28) organotypic slices usually lacked CR+ cells; immunostained neurons were mainly present in patches (Fig. 7A). Cajal-Retzius cells were preserved all over the culturing period, even in slices fixed at D35. The morphology of CR+ interneurons has also changed in the organotypic cultures. Cells with small soma usually disappeared earlier than neurons with a large cell body. Furthermore, CR+ neurons with no or very short dendrites were also frequently observed. In the second half of the culturing period (at D21-42), cells with beaded or distorted dendrites were also often detected.

**Figure 7:**
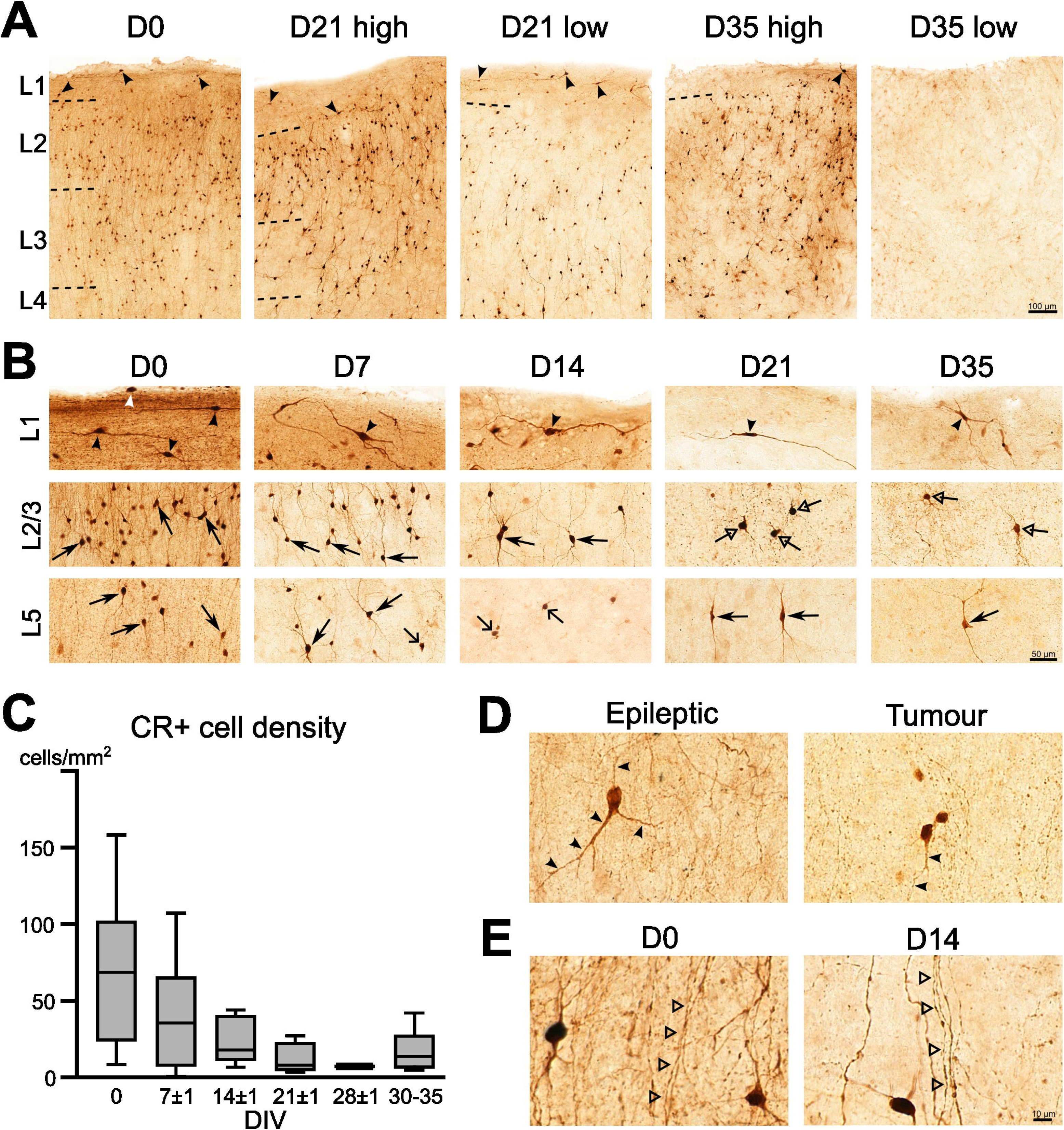
The number of CR+ cells considerably decreases during the culturing period. **(A)** Calretinin-stained sections from acute sample (D0) and organotypic cultures (D21, D35). Numerous CR+ inhibitory interneurons are present in the human neocortex. Large horizontal bipolar cells are identified as Cajal-Retzius cells in layer 1 (arrowheads, L1). CR-stained neurons with high and low density are shown from organotypic slices fixed at D21 and D35. The images showing D21 are from different slices but from the same HOC sample (HOC25), whereas the images representing D35 are from the same slice. The neocortical layers cannot be separated at the later stages of the culture period. **(B)** Images show CR+ neurons in the different layers of the human neocortex at different time points during the culturing period. Cajal-Retzius cells (arrowheads) are present in L1, from D0 to D35. Note the large cell body and the horizontal dendrites. Numerous cells are visible in L2/3 and L5 at D0, whereas their density decreases with culturing time. Arrows point on typical bipolar and multipolar cells, open arrows show the CR+ neurons with small cell body and very short dendrites, and open triangle arrows point on CR+ cells with distorted dendrites. These latter cells were usually seen at the later stages of the culture period. **(C)** The density of CR+ neurons from D0 to D35 is decreasing. Statistical tests did not show significant differences between the examined days of the week-by-week analysis. **(D)** CR-positive inhibitory cells usually had a more complex dendritic tree in epileptic (left image) than in non-epileptic tumour (right image) tissue. Arrowheads point on the dendrites of the CR+ neuron. **(E)** Parallelly running CR+ dendrites were frequently seen in acute samples (D0), as well as at the earlier stages of the culturing time. Open triangle arrows point to the dendrite bundles.

Viral treatment affected both the cell density and the morphology of the CR+ neurons. In most cases, the number of cells decreased considerably in virus-injected slices compared to non-injected slices from the same HOC sample. In virally treated slices we found large areas lacking CR+ neurons, and few spots with CR+ cells which possessed either very short and/or distorted, beaded dendrites. Similar observations were made when comparing slices kept in hCSF vs. AM. Fewer cells were visible in hCSF than in AM, and they had more pathological morphology. Larger areas devoid of CR+ neurons were more typical to hCSF and virally treated slices than to AM or non-injected ones.

To assess the changes in the cell density of CR+ interneurons, we performed semi-automatic cell counting (see Methods) on the above mentioned 43+11 slices (for values see Table 1 and Supplementary Table 9). Although the morphology of the CR+ cells in the epileptic and tumour tissue was usually different, we did not find significant differences between epileptic and tumour patients (Mann-Whitney test, *p*=0.1143), therefore these samples were pooled. No significant differences could be shown when comparing tissues derived from different patients (HOC comparison, Krukal-Wallis ANOVA, *p*=0.2865). The density of CR+ cells did not differ when examined with a day-by-day analysis (Kruskal-Wallis ANOVA, *p*=0.1743). Despite the visible decrease in cell density over the culturing period, the week-by-week changes were significantly different only in general (all data pooled together, ANOVA, *p*=0.0182), but the post-hoc test did not show any significant difference between the examined periods. Despite the morphological changes and the visible reduction in the cell density across different media and viral treatments, neither affected significantly cell density (CSF vs. AM: Mann-Whitney test, *p*=0.1624; viral treatment vs. control in AM: paired t-test, *p*=0.3447).

## Discussion

### Functional readouts: electrophysiology and calcium imaging

Human organotypic slice cultures preserved their ability to generate spontaneous population activity detected with both electrophysiology and Ca^2+^-imaging. The spontaneous population events observed here lacked the hallmarks of epileptiform discharges reported in organotypic cultures under pathological conditions: the highly stereotyped, large-amplitude bursts with near-synchronous recruitment of neuronal populations^3,25^. In contrast, the events detected in the present study occurred within physiological firing range and displayed heterogeneous recruitment across neuronal classes, consistent with coordinated population dynamics rather than pathological hypersynchrony^12,13^.

The observed cellular composition was physiologically plausible and consistent across patients and time points, with principal cells comprising approximately 75–80% of all recorded single neurons. This proportion closely matches pyramidal-to-interneuron ratios reported in rodent and human cortical and hippocampal circuits in organotypic preparations^4,5^. Reliable separation of interneurons and principal cells based on spike waveform further confirms that canonical electrophysiological cell type identities are retained in culture^26^ and principal neurons remain the dominant contributors ot spontaneous network activity in vitro^13,27^.

Spike waveform dynamics revealed a significant narrowing of action potential half-width in principal cells over the culture period, while interneuron waveforms remained comparatively stable. Experimental and modeling studies indicate that activity-dependent maturation processes can modulate sodium channel density and spike initiation dynamics, leading to sharper action potentials and improved temporal precision^28,29^. Because fast-spiking interneurons already operate with optimized spike kinetics, their extracellular waveforms may be less susceptible to such plasticity. The selective evolution of pyramidal-cell spike kinetics therefore likely reflects adaptive intrinsic plasticity during circuit reorganization in vitro.

Several electrophysiological metrics followed coherent temporal trajectories during the culture period. Burst generation in principal cells increased, and discharge irregularity (ISI CV) transiently increased during the intermediate culture phase (D7–15) before returning toward baseline values by D22. This sequence closely resembles the well-described phases of post-slicing network adaptation: initial deafferentation, transient hyperexcitability driven by reactive synaptogenesis, and subsequent homeostatic stabilization of circuit activity^30,31^. Comparable dynamics have been reported in both rodent and human slice preparations, where early circuit perturbations are gradually compensated by intrinsic and synaptic homeostatic mechanisms^31,32^.

Analysis of single-unit recruitment during spontaneous population activity events further revealed distinct cell-type-specific participation patterns. Intrinsically bursting principal cells were the most consistently recruited neuronal class (93.75%), supporting that burst-capable pyramidal neurons contribute disproportionately to the generation of synchronous population activity^13,33^. Interneurons were recruited less frequently overall, but those that participated in population events exhibited approximately twofold higher firing rates than non-recruited interneurons, suggesting that strongly driven and highly active inhibitory neurons also participate in the generation and maintenance of population activity. This is consistent with the rapid engagement of fast-spiking interneurons during synchronized cortical activity and their role in shaping network timing and gain control^5,34^.

SPA properties showed no statistically significant change across recording days, indicating that the capacity for generating population activity is maintained throughout the culture period. However, several electrophysiological features evolved coherently over culturing time. Multiple unit activity, and single-unit level features such as discharge regularity (ISI CV) and the proportion of recruited neurons reflected a transient perturbation during the early culture phase (D7-15), then returned toward D0 values by D22. These findings suggest that, following an early phase of network disturbance – presumably linked to the considerable neuron loss – human cortical organotypic cultures reach a state by around three weeks in which single-unit firing patterns and population-event recruitment more closely resemble those of the acute tissue. This contrasts with the progressive development reported in rodent organotypic cultures, where developmental stage and genetic background of the original brain tissue strongly influence network maturation^35^. In human preparations using adult tissue, however, intrinsic circuit dynamics reorganize towards reproducible neuronal activity patterns during prolonged in vitro maintenance^7^. Our findings extend this observation by demonstrating that both single-unit firing and population recruitment patterns stabilize within approximately three weeks, defining a reproducible window for experimental functional investigation.

Wide-field Ca^2+^-imaging provided complementary evidence for functional stabilization by following the same slices repeatedly over the culturing period. Recurrent population Ca²C transients were detected in line with the development of GCaMP-expression, which are most probably the optical readouts of electrophysiologically recorded SPAs. This activity persisted for weeks, forming spatially confined and temporally stable activity hubs. These active regions were consistently observed across repeated recordings, indicating that network organization is not random but constrained by underlying structural connectivity. The reliable appearance of the synchronous Ca^2+^-transients indicates that the functional outputs of the human cortex are preserved in culture. Similar stable activity patterns have been reported in human hippocampal slice cultures and are thought to reflect preserved microcircuit motifs^7,8^. The persistence of these population activities, together with the convergence of single-unit dynamics, suggests that network-level attractor states emerge despite ongoing neuronal loss and structural remodeling.

### Histology

Histological analysis revealed substantial structural reorganization in human cortical organotypic cultures. As in rodent organotypic cultures^36^ neuronal density decreased significantly in our human slice cultures, particularly during the first week, accompanied by the partial degradation of the laminar architecture^11,37^. Neuronal loss was very variable across different samples, but also at the temporal and spatial dimensions. Remarkably different neuronal density could be detected at the same time point of the culturing period, as well as in different slices from the same sample (even at the same day in vitro). Neuron loss could affect the entire neocortex, or was confined to specific layers or a specific area, especially related to viral injection. This spatial/temporal variability might be related to the diversity of the patients, and thus might be a human-specific feature. However, operation and slicing circumstances can also affect diverse neuronal survival^4^.

Inhibitory interneuron populations were particularly vulnerable. With a similar timeline to the overall neuronal loss, the density of both parvalbumin- and calretinin-positive interneurons declined over time, with marked regional loss and morphological modification. However, the surviving parvalbumin- and calretinin-positive cells preserved their functional identity: both types of perisomatic interneurons (PV+ axo-axonic and basket cells) as well as CR+ Cajal-Retzius cells were observed even at the latest stages of the culturing period. The loss of these cell types is consistent with prior reports highlighting the susceptibility of inhibitory neurons to stress, such as human epilepsy^38,39^ or ischaemia^40^. The survival of Cajal-Retzius cells is in contrast with findings in mouse organotypic cultures, where this cell type - together with all CR+ inhibitory cells - are totally lost during the first week^41^. However, this might be a human-specific feature, as unlike in rodents^42^, Cajal-Retzius cells persist in the adult human neocortex^43^.

Glial responses were prominent, quickly developed and persisted until the end of the culturing period. Similar to the neuronal loss, the degree of both astroglial and microglial coverage was variable, patches containing a sparse glial network intermingled with highly covered areas. Astrocytes exhibited a pronounced reactive gliosis, particularly at the slice surfaces, consistent with injury-induced astrocytic reorganization (for review see ^44^). Microglial cells displayed diverse activation states, transitioning from homeostatic to reactive and phagocytic phenotypes, in line with their roles in synaptic remodeling and debris clearance^45^. Notably, while glial morphology changed dramatically, the overall coverage remained relatively stable, suggesting that qualitative rather than quantitative changes in glial function accompany network reorganization.

In summary, substantial neuronal loss involving both excitatory principal cells and inhibitory interneurons are accompanied by massive gliosis. Both astroglial and microglial cells show strikingly activated morphology and develop an extensive glial sheet on the cut surfaces of the slices. Despite these changes, functional activity persisted, indicating that network function is maintained within a structurally evolving substrate, and the surviving minority drive the network dynamics.

## Conclusions

Conceptually, our findings indicate that structural degeneration and functional stabilization are not opposing processes but co-emergent features of network adaptation. While neuronal loss, gliosis, and interneuron vulnerability reshape the anatomical organisation, intrinsic and synaptic homeostatic mechanisms drive the network toward stable attractor states that preserve coherent activity patterns. The combination of electrophysiology, longitudinal Ca^2+^-imaging, and detained quantitative histology demonstrates that human organotypic cultures do not simply deteriorate over time but instead reorganize into reproducible functional systems. Human slice cultures advance from an early perturbed state toward a reorganized system in which key features of its output functions — including population synchrony, cell-type-specific firing patterns, and recruitment dynamics — are maintained. This integrated perspective highlights the value of human organotypic cultures as a translational platform: despite profound perturbation and ongoing structural remodeling, they retain key principles of human circuit organization while offering experimental accessibility unavailable in vivo.

## Data availability

Data (including electrophysiology analysis software) are available upon request from the corresponding author (L.W.).

## Supplementary material

Supplementary material is available at *Brain* online.

## Author contributions

**Tóth, Estilla Zsófia: Conceptualization, Data curation, Investigation, Methodology, Resources, Validation**

Made all organotypic cultures up to HOC29, lead all immunostainings (including recutting the slices) and all wide-field imaging. Lead the scanning process of all stained sections.

**Stelcz, Rebeka: Conceptualization, Data curation, Formal analysis, Investigation, Methodology, Resources, Software, Visualization, Writing – original draft**

Helped in the culturing from the beginning, continued it alone from HOC30. Participated in the recutting, staining, scanning, lead the entire quantitative histology part with statistics. Made NeuN, GFAP, IBA1 detection and quantitative analysis alone, lead PV and CR cell counting. Wrote the paper, made figures.

**Bod, Réka: Conceptualization, Data curation, Formal analysis, Methodology, Software, Writing – original draft**

Developed the framework for the ephys analysis pipeline, wrote the SPA detecting and clustering software. Clustered all single cells, analysed the ephys SPA part, made all statistics for the ephys part. Participated in the human operations. Wrote the paper.

**Petik, Ábel: Conceptualization, Data curation, Formal analysis, Investigation, Methodology, Software, Visualization.**

Developed the custom wide-field imaging platform and software pipeline for longitudinal calcium imaging; performed and analysed wide-field imaging experiments.

**Juhász, Eszter: Data curation, Formal analysis, Investigation, Resources**

Cell counting (half-manual) for PV and CR sections, participated in human tissue preparation, recutting and staining the slices, part of human data administration.

**Tóth, Kinga: Conceptualization, Data curation, Investigation, Methodology, Project administration, Resources, Supervision, Validation, Writing – original draft**

Lead and performed most ephys experiments, lead SPA detection, wrote the paper, made human tissue preparation, lead and made the administration of human data

**Szalai, Liza: Data curation, Investigation**

Recutting the slices, staining, scanning and analysis of GFAP+ sections

**Essam, Nour: Investigation, Project administration**

Performed ephys experiments, administration of ephys experiments, participated in human tissue preparation

**Somogyi, Fanni: Data curation, Investigation**

Performed wide-field imaging experiments.

**Kovács, Beatrix: Methodology, Resources**

Provided viral tools used for imaging experiments.

**Mojtahedzadeh, Amirmahdi: Data curation**

Made most SPA detection.

**Bagó, Attila György: Project administration, Resources**

Neurosurgeon, provided human tissue.

**Erőss, Loránd: Project administration, Resources**

Neurosurgeon, provided human tissue.

**Orbay, Péter: Project administration, Resources**

Neurosurgeon, provided human tissue.

**Szabó, Johanna Petra: Project administration, Resources**

Organized the operations, helped in providing clinical data.

**Fabó, Dániel: Project administration, Resources**

Neurologist, provided human clinical data.

**Hajnal, Boglárka: Project administration, Resources**

Neurologist, provided human clinical data.

**Rácz, Bence: Resources, Supervision, Writing – review & editing**

Provided scanning possibilities, taught scanning method, reviewed and edited the manuscript.

**Hillier, Daniel: Conceptualization, Funding acquisition, Methodology, Resources, Supervision, Software, Writing – review & editing**

Conceived the longitudinal wide-field imaging approach, supervised the development of the imaging platform and associated analysis workflow, and contributed to the viral strategy/tools used for the imaging experiments; reviewed and edited the manuscript.

**Ulbert, István: Conceptualization, Funding acquisition, Methodology, Project administration, Software, Writing – review & editing**

Concept of ephys recordings and analysis (both SPA detection and clustering), supervision of the ephys pipeline development and analysis, wrote ethical approval, reviewed and edited the manuscript.

**Wittner, Lucia: Conceptualization, Data curation, Funding acquisition, Investigation, Methodology, Project administration, Resources, Supervision, Visualization, Writing – original draft**

Concept of the overall study, coordination of the experimental workflow, lead and participated in human tissue preparation, contributed to the ephys pipeline development, supervision of the entire histology and ephys parts, wrote ethical approval, wrote the paper, made figures.

