## Supplementary Material for "Preserved functions with profound morphological reorganization in human organotypic cultures"

1) Patients and tissue samples

Postoperative tissue samples derived from 30 patients were used for preparing cortical organotypic slice cultures. Supplementary Table 1 shows patient data.

**Supplementary Table 1. Patient data.**

F=female, M=male, NoEpi: tumor patient without epileptic seizure, NoMed: tumor patient with one epileptic seizure, and no need for antiepileptic medication, TreatEpi: epileptic patient, seizure free with appropriate medication, is operated because of a tumor; ResEpi: therapy resistant epileptic patient; close from tumor: < 3 cm, distant: > 3 cm.

| Patient ID | Age | Gender | Stage of epilepsy | Duration of epilepsy | Resected cortical region | Distance from tumor | Histology/Diagnosis |
| --- | --- | --- | --- | --- | --- | --- | --- |
| HOC 1 | 70 | F | NoEpi | - | frontal | close | glioblastoma grade 4 |
| HOC 2 | 65 | F | NoEpi | - | parietal | close | glioblastoma grade 4 |
| HOC 3 | 66 | F | NoEpi | - | frontal | close | lung adenocarcinoma metastasis |
| HOC 4 | 68 | M | TreatEpi | N/A | temporal | close | glioblastoma grade 4 |
| HOC 5 | 82 | F | NoEpi | - | temporal | close | lung adenocarcinoma metastasis |
| HOC 6 | 43 | M | TreatEpi | 2 years | temporal | close | astrocytoma grade 3 |
| HOC 7 | 61 | M | TreatEpi | 1 month | temporal | close | neuroendocrin carcinoma metastasis |
| HOC 8 | 42 | M | ResEpi | 38 years | temporal | - | hippocampal sclerosis |
| HOC 9 | 27 | M | ResEpi | 7 years | temporal | - | hippocampal sclerosis |
| HOC 10 | 24 | M | TreatEpi | 4 years | temporal | close | astrocytoma grade 3 |
| HOC 11 | 58 | F | NoEpi | - | frontal | close | glioblastoma grade 4 |
| HOC 12 | 76 | M | NoEpi | - | temporal | N/A | lung adenocarcinoma metastasis |
| HOC 13 | 58 | F | NoEpi | - | temporal | close | lung adenocarcinoma metastasis |
| HOC 14 | 62 | M | TreatEpi | 2 weeks | frontal | close | pulmonary squamous cell carcinoma metastasis |
| HOC 15 | 79 | M | NoEpi | - | parietal | close | adenocarcinoma metastasis |
| HOC 16 | 67 | M | NoEpi | - | temporal | close | glioblastoma grade 4 |
| HOC 17 | 74 | M | NoMed | 1 month | temporal | close | glioblastoma grade 4 |
| HOC 18 | 68 | F | TreatEpi | 2 weeks | temporal | close | glioblastoma grade 4 |
| HOC 19 | 51 | M | TreatEpi | 2 weeks | frontal | N/A | glioblastoma grade 4 |
| HOC 20 | 60 | M | NoEpi | - | temporal | close | glioblastoma grade 4 |
| HOC 21 | 58 | M | NoEpi | - | temporal | close | adenoid glioblastoma grade 4 + meningeoma grade 1 |
| HOC 23 | 36 | M | ResEpi | 14 years | temporal | - | FCD 1A + gliosis |
| HOC 24 | 48 | F | ResEpi | 34 years | temporal | - | hippocampal sclerosis |
| HOC 25 | 23 | M | ResEpi | 11 years | temporal | - | gliosis + microglia activation |
| HOC 26 | 75 | M | NoEpi | - | temporal | distant | glioblastoma grade 4 |
| HOC 27 | 56 | M | NoEpi | - | temporal | close | small cell neuroendocrine carcinoma metastasis |
| HOC 28 | 55 | M | NoEpi | - | temporal | close | glioblastoma grade 4 |
| HOC 29 | 44 | M | ResEpi | 8 years | temporal | distant | cavernoma + gliosis |
| HOC 30 | 35 | F | ResEpi | 5 years | temporal | - | hippocampal sclerosis |
| HOC 35 | 39 | F | ResEpi | 16 years | temporal | distant | dysembryoplastic neuroepithelial tumor |

Supplementary Table 2. shows the number of organotypic slices made from the different tissue samples, their culturing circumstances, as well as their use for different experiments, such as viral injection, wide-field imaging, electrophysiology and anatomical examinations.

**Supplementary Table 2. Number of organotypic slices used for different experiments.**

Note, that tissue samples used and/or fixed at the day of the operation (D0) are in parenthesis, and are not included in the number of organotypic slices (column of Total slices), as these slices were not cultured. HOC: human organotypic culture from one patient, see also in Suppl. Table 1; hCSF: human cerebrospinal fluid, AM: artificial medium, PS: penicillin+streptomycin, PSA: penicillin+streptomycin+amphotericinB, Ephys: electrophysiological recording, NeuN: neuronal nuclear marker; GFAP: glial fibrillar acidic protein, astroglial marker; IBA1: microglia marker; PV: parvalbumin, CR: calretinin. Note, that usually several stainings were made from each slice, and those stainings were not necessarily used for the anatomical investigation.

| **HOC** | **Total slices** | **hCSF** | | **AM (PSA)** | **Viral treated** | **Widefield** | **Ephys** | **Stained** | **NeuN** | **GFAP** | **IBA1** | **PV** | **CR** |
| --- | --- | --- | --- | --- | --- | --- | --- | --- | --- | --- | --- | --- | --- |
|  |  | **PS** | **PSA** |  |  |  |  |  |  |  |  |  |  |
| 1 | 7 | 7 |  |  | 0 |  | 0 (0) | 6 | 0 (0) | 4 (0) | 0 (0) | 0 (0) | 0 (0) |
| 2 | 17 | 17 |  |  | 0 |  | 0 (0) | 2 | 0 (0) | 1 (1) | 0 (0) | 0 (0) | 0 (0) |
| 3 | 19 | 19 |  |  | 0 |  | 0 (0) | 3 | 0 (0) | 2 (0) | 0 (0) | 0 (0) | 0 (0) |
| 4 | 10 | 10 |  |  | 0 |  | 0 (0) | 5 | 4 (0) | 4 (1) | 0 (0) | 2 (0) | 0 (0) |
| 5 | 9 | 9 |  |  | 0 |  | 0 (0) | 1 | 1 (0) | 1 (1) | 0 (0) | 0 (0) | 0 (0) |
| 6 | 16 | 16 |  |  | 0 |  | 0 (0) | 11 | 8 (1) | 6 (2) | 0 (0) | 4 (1) | 0 (0) |
| 7 | 8 | 8 |  |  | 0 |  | 0 (0) | 6 | 4 (1) | 4 (1) | 2 (0) | 1 (1) | 0 (1) |
| 8 | 12 |  | 12 |  | 0 |  | 0 (0) | 5 | 3 (1) | 3 (1) | 3 (1) | 0 (1) | 0 (0) |
| 9 | 20 |  | 20 |  | 0 |  | 0 (0) | 14 | 12 (2) | 11 (2) | 5 (2) | 0 (1) | 0 (0) |
| 10 | 11 |  | 11 |  | 0 |  | 0 (0) | 6 | 5 (0) | 4 (0) | 0 (0) | 1 (0) | 3 (0) |
| 11 | 7 |  | 7 |  | 0 |  | 0 (0) | 5 | 4 (1) | 0 (0) | 0 (0) | 1 (1) | 1 (1) |
| 12 | 6 |  |  | 6 | 0 |  | 1 (4) | 1 | 1 (0) | 0 (0) | 0 (0) | 1 (0) | 1 (0) |
| 13 | 8 |  | 4 | 4 | 5 | 6 | 0 (6) | 8 | 8 (0) | 5 (0) | 1 (0) | 5 (0) | 6 (0) |
| 14 | 3 |  |  | 3 | 2 |  | 0 (1) | 3 | 2 (0) | 2 (0) | 0 (0) | 1 (0) | 0 (0) |
| 15 | 5 |  |  | 5 | 4 | 5 | 1 (4) | 5 | 3 (0) | 0 (0) | 0 (0) | 4 (0) | 0 (0) |
| 16 | 4 |  |  | 4 | 2 |  | 0 (3) | 3 | 3 (0) | 2 (0) | 0 (0) | 0 (0) | 0 (0) |
| 17 | 9 |  |  | 9 | 2 |  | 0 (0) | 5 | 5 (1) | 1 (1) | 0 (0) | 3 (1) | 1 (1) |
| 18 | 14 |  |  | 14 | 10 |  | 1 (0) | 12 | 7 (1) | 11 (1) | 0 (0) | 2 (0) | 4 (0) |
| 19 | 4 |  |  | 4 | 0 |  | 0 (1) | 0 | 0 (1) | 0 (1) | 0 (1) | 0 (1) | 0 (1) |
| 20 | 3 |  |  | 3 | 3 |  | 0 (3) | 3 | 0 (1) | 3 (1) | 0 (0) | 1 (0) | 0 (0) |
| 21 | 16 |  |  | 16 | 6 | 6 | 1 (0) | 15 | 13 (1) | 5 (1) | 3 (0) | 4 (1) | 2 (1) |
| 23 | 24 |  |  | 24 | 7 |  | 0 (0) | 10 | 10 (0) | 10 (0) | 7 (0) | 8 (0) | 6 (0) |
| 24 | 8 |  |  | 8 | 0 |  | 0 (6) | 6 | 6 (1) | 6 (1) | 6 (1) | 6 (1) | 5 (1) |
| 25 | 4 |  |  | 4 | 0 |  | 2 (6) | 2 | 1 (1) | 1 (1) | 1 (1) | 2 (1) | 1 (1) |
| 26 | 9 |  |  | 7 | 0 |  | 3 (6) | 6 | 6 (1) | 6 (1) | 5 (1) | 4 (1) | 4 (1) |
| 27 | 5 |  |  | 5 | 0 |  | 3 (2) | 5 | 5 (1) | 5 (1) | 3 (1) | 3 (1) | 4 (1) |
| 28 | 5 |  |  | 5 | 0 |  | 2 (3) | 3 | 3(1) | 3 (1) | 2 (1) | 3 (1) | 2 (1) |
| 29 | 6 |  |  | 6 | 1 |  | 4 (5) | 4 | 4 (1) | 4 (1) | 2 (1) | 3 (1) | 3 (1) |
| 30 | 15 |  |  | 15 | 0 |  | 5 (6) | 0 | 0 (0) | 0 (0) | 0 (0) | 0 (0) | 0 (0) |
| 35 | 13 |  |  | 13 | 0 |  | 5 (6) | 0 | 0 (0) | 0 (0) | 0 (0) | 0 (0) | 0 (0) |
|  | **297** | **86** | **54** | **155** | **42** | **15 (2)** | **28 (62)** | **155** | **118 (17)** | **104 (20)** | **40 (10)** | **59 (14)** | **43 (11)** |

**In vitro electrophysiology**

**Population activity**

Cellular firing as well as the presence of spontaneous population activity12 (SPA) emerging in standard physiological baths were noted in all slices (Fig. 1). In the acute samples (D0) cell firing could be detected in 73% of the slices (45/62), 37% of the slices (23/62) generated SPA as well. In the case of D7 organotypic slices 56% (5/9) and 44% (4/9) displayed cellular firing and SPA, respectively. Neuronal firing was observed in 100% (5/5) of the D14 organotypic samples, of which 60% (3/5) also produced SPA. In the case of D21 organotypic slices 78% (7/9) and 33% (3/9) displayed cellular firing and SPA, respectively. In the D30 organotypic tissues neuronal firing could be detected in 100% of the slices (4/4) and 50% of the slices (2/4) generated SPA as well, and finally the D35 slice showed cell firing without the emergence of SPA.

We examined the basic network properties of SPAs generated by different slices both in acute (D0) and in cultured slices. Suppl. Table 3 shows the properties of SPAs along the culturing period, at D0, D7, D15, D22 and D30, such as recurrence frequency events, ISI mean (ISI = inter event interval in this case), average LFPg, CSD and MUA amplitudes.

**Cell-type classification and sample composition**

A total of 151 single units (neurons) were recorded across four patients (HP108, HP110, HP112, HP114) and four time points (D0, D7, D15, D20; we excluded D30 from all group analyses due to a single available recording). Units were classified into three fine cell-type categories: regularly spiking principal cells (RS-PC, n = 99), intrinsically bursting principal cells (IB-PC, n = 16), and interneurons (IN, n = 36). For coarse-level comparisons, RS-PC and IB-PC were pooled as “excitatory” principal cells (n = 115) and contrasted with “inhibitory” interneurons (n = 36). The proportion of principal cells relative to interneurons was broadly consistent across patients (HP108: 50% PC; HP110: 75%; HP112: 77.1%; HP114: 82.8%) and across days-in-vitro (D0: 78.1%; D7: 73.9%; D15: 69.2%; D20: 82.1%), with IB-PCs representing a minority subtype that became more prevalent at later time points (D0: 9.6%; D20: 25.0%).

**Electrophysiological properties by cell type**

We defined several cellular properties (metrics) for each clustered cell, such as the average firing rate (Hz), the maximum firing rate over the most active 10 s window (Hz), the burstiness 1 (% of spikes in bursts, see Methods for definition of burst), the burstiness 2. indexed by inter-spike intervals shorter than 20 ms (%), spike half-width 1 (ms), spike half-width 2 (ms), and inter-spike interval coefficient of variation (ISI CV). Descriptive statistics for all metrics are provided in Suppl. Tables 4-7 (fine, coarse and overall cell types).

**Firing rate and maximum firing rate**. Mean firing rates were comparable across RS-PC (1.66 ± 2.11 Hz), IB-PC (1.11 ± 0.71 Hz), and IN (1.47 ± 1.73 Hz; Kruskal–Wallis: H = 0.26, p = 0.877, ns). Differences in the maximum firing rates over the most active 10 s window were similarly non-significant across fine cell types (H = 0.37, p = 0.833, ns) and between principal cells and interneurons (H = 0.37, p = 0.545, ns), indicating that gross firing rate did not reliably distinguish cell-type classes in this preparation.

**Burstiness.** Burst metrics showed strong and significant cell-type dependence. As expected, IB-PCs displayed markedly higher burstiness by percentage (3.72 ± 3.74%) and higher burstiness indexed by ISI<20 ms (35.4 ± 14.1%) compared with RS-PCs (0.37 ± 1.08%; 13.1 ± 13.6%) and INs (0.24 ± 0.85%; 17.5 ± 25.4%). Kruskal–Wallis tests confirmed significant differences for both measures across fine cell types (burstiness%: H = 36.13, p<0.001; burstiness ISI< 20 ms: H = 22.00, p<0.001). Pairwise comparisons (Bonferroni-corrected) revealed that IB-PCs differed significantly from both RS-PCs (burstiness%: p<0.001; ISI<20 ms burstiness: p<0.001) and INs (burstiness%: p<0.001; ISI<20 ms burstiness: p = 0.006), while RS-PCs and INs did not differ on either measure. At the coarse level, principal cells exhibited significantly higher burstiness than interneurons (burstiness%: H = 6.05, p = 0.014; coarse pairwise p = 0.014), though the ISI < 20 ms index did not reach significance after correction (H = 2.84, p = 0.092).

**Spike waveform metrics**. Half-width 1 (the main peak/trough half-width) differed significantly across fine cell types (H = 72.21, p < 0.001), with INs showing substantially narrower waveforms (0.178 ± 0.032 ms) compared with RS-PCs (0.275 ± 0.066 ms; p < 0.001) and IB-PCs (0.302 ± 0.053 ms; p < 0.001), consistent with the fast-spiking physiology of interneurons. RS-PCs and IB-PCs did not differ significantly in half-width 1 (p = 0.147). Half-width 2 (secondary peaks’ values) showed the same pattern (IN: 0.354 ± 0.185 ms; RS-PC: 0.577 ± 0.193 ms; IB-PC: 0.638 ± 0.118 ms; H = 31.73, p < 0.001), with significant separation between INs and both principal cell subtypes (p < 0.001 for both), but not between RS-PC and IB-PC (p = 0.636). At the coarse level, principal cells had significantly wider waveforms than interneurons for both half-width metrics (half-width 1: H = 69.79, p < 0.001; half-width 2: H = 30.28, p < 0.001).

**ISI coefficient of variation**. ISI CV did not differ across fine cell types (RS-PC: 1.79 ± 1.16; IB-PC: 1.68 ± 0.91; IN: 2.12 ± 1.62; H = 0.38, p = 0.827, ns) or between principal cells and interneurons (H = 0.30, p = 0.585, ns), indicating that discharge regularity was not systematically related to morphological cell-type classification in these cultures.

**Electrophysiological properties across days *in vitro***

We examined the electrophysiological properties of neurons detected on the different time points of the culturing period. For this analysis, we excluded the single neuron clustered in the recording made on D30. To see the effects of culturing, we also pooled all culture days (D>0: D7, D15, D22 combined) and compared to D0.

*Firing rate.* A significant days-in-vitro (DIV) effect was detected (KW: H = 17.70, p < 0.001), with D15 showing the lowest mean firing rate (0.60 ± 0.72 Hz) compared with D0 (1.71 ± 2.03 Hz; p < 0.001) and D7 (2.11 ± 2.26 Hz; p = 0.004). D22 (1.64 ± 1.89 Hz) was not significantly different from D0.

*Burstiness ISI < 20 ms.* This metric increased progressively across DIV (D0: 11.5 ± 12.9%; D7: 8.86 ± 15.7%; D15: 32.2 ± 24.2%; D22: 20.8 ± 17.3%; KW: H = 25.16, p < 0.001), reflecting a marked increase in high-frequency bursting at D15 compared with D0 (p < 0.001) and D7 (p = 0.002), with D20 also differing from D7 (p = 0.036).

*Spike half-width 1.* Waveforms narrowed over time (D0: 0.276 ± 0.082 ms; D15: 0.214 ± 0.044 ms; KW: H = 14.53, p = 0.002), with significant difference between D0 and D15 (p = 0.002). Narrowing of the action potential upstroke is consistent with the maturation of voltage-gated sodium channel expression in maturing networks^1^.

*ISI CV.* The most pronounced DIV-dependent effect was seen for discharge irregularity (KW: H = 28.60, p < 0.001). ISI CV was lowest at D0 (1.46 ± 0.76) and D20 (1.47 ± 0.85), and highest at D15 (2.70 ± 1.32) and D7 (2.59 ± 1.99). Pairwise comparisons showed significant differences between D0 and D15 (p < 0.001) and between D15 and D20 (p < 0.001). Maximum firing rate, burstiness %, and half-width 2 did not show significant DIV-dependent effects.

**Acute versus culture changes (D0 vs D>0)**

To assess the effects of the culturing on the human cortical tissue, units recorded at D0 (n = 73) were compared with all units recorded at subsequent time points combined (D>0; n = 77). Across all cells pooled, significant increases in burstiness ISI < 20 ms (p = 0.044), half-width 1 narrowing (p = 0.002), and ISI CV (p < 0.001) were observed from D0 to D>0. When analyzed by cell type, RS-PCs showed a significant increase in firing rate at D>0 relative to D0 (p = 0.033), increased ISI CV (p = 0.003), and narrower half-width 1 (p < 0.001). IB-PCs showed a significant increase in burstiness % between D0 and D>0 (p = 0.020). Interneurons showed a marked and significant increase in ISI CV (p = 0.003), indicating progressively more irregular discharge patterns over the culture period. No significant D0 vs D>0 differences were found for maximum firing rate in any cell-type subgroup.

**Event-responsiveness and its relationship to electrophysiological properties**

We determined for each cell whether it increases its firing rate during spontaneous population activity with an adapted Monte Carlo approach (see Methods, ^2^). Cells with increased firing rate were considered as responsive neurons.

*Proportional analysis.* The proportion of neurons with increased firing rate differed significantly across fine cell types (chi-squared test: χ² = 9.55, p = 0.008): IB-PCs showed the highest proportion of responsive neurons (93.75%, 15/16), followed by RS-PCs (66.67%, 66/99) and INs (50.0%, 18/36). At the coarse level, a significant difference was also observed between principal cells and interneurons (70.43% vs 50.0%; χ² = 4.21, p = 0.040). The proportion of responsive cells varied significantly across DIVs (D0: 71.2%; D7: 39.1%; D15: 65.4%; D20: 71.4%; χ² = 8.55, p = 0.036), with D7 showing a notably lower proportion of responsive cells. No significant inter-patient difference was found (p = 0.071).

*Metric differences between responsive and non-responsive neurons.* For RS-PCs, responsive cells showed significantly higher burstiness ISI < 20 ms than non-responsive cells (mean 15.33 ± 13.44% vs 8.76 ± 13.19%; Mann–Whitney p < 0.001, rank-biserial r = −0.44), indicating that event-responsive RS-PCs engaged more high-frequency burst firing during population events. No other metric reached significance for RS-PCs after correction. For INs, responsive neurons had significantly higher firing rates (2.20 ± 2.13 Hz vs 0.75 ± 0.70 Hz; p = 0.022, r = −0.45) and lower burstiness % (0.0 ± 0.0% vs 0.48 ± 1.17%; p = 0.040, r = 0.22), suggesting that event-responsive interneurons tended to be tonically active, regularly firing, most probably fast-spiking cells. For pooled principal cells, increased cells showed significantly higher burstiness % (p = 0.030, r = −0.22) and higher burstiness ISI < 20 ms (18.69 ± 14.94% vs 10.38 ± 16.07%; p < 0.001, r = −0.46), a pattern consistent across the RS-PC and coarse-level neuron subgroups. IB-PC subgroup comparisons were underpowered due to only a single non-responsive cell (n_indifferent = 1) and should be interpreted with caution.

In summary, we can conclude, that IB-PC tend to participate more in SPA events than other neuron types, responsive RS-PCs have higher burstiness, than those with unchanged firing rate. We can speculate whether these RS-PCs participate in the generation of SPA events because of their higher burstiness, or their burstiness is higher because they can be better recruited in population events. Furthermore, the fact, that responsive inhibitory interneurons had higher firing rate suggests that fast-spiking cells are more participating in SPA events than other interneuron subtypes.

**Supplementary Table 3. Network properties of SPAs.** Mean ± SD for the network properties of SPAs, such as the recurrence frequency, the ISI mean (showing the mean interval between SPA events), LFPg, CSD and MUA amplitude (see Methods). None of the examined properties showed significant differences between the different recording days. D7: n=3 SPAs from 3 slices, 3 patients; D15: n=4 SPAs in 3 slices in 2 patients; D22: n=4 SPAs in 3 slices in 2 patients and D30: n=1 SPA in 1 slice from 1 patient. KW: Kruskal-Wallis ANOVA, Sig: significance, column with green shading shows values for all SPA pooled.

| **N recordings** | **D0 (n=11)** | **D7 (n=3)** | **D15 (n=4)** | **D22 (n=4)** | **D30 (n=1)** | **All (n=23)** | **KW p** | **Sig** |
| --- | --- | --- | --- | --- | --- | --- | --- | --- |
|  | **Mean ± SD** | **Mean ± SD** | **Mean ± SD** | **Mean ± SD** | **Value** | **Mean ± SD** |  |  |
| **N events** | 183 ± 184 | 94.3 ± 135 | 61 ± 24.9 | 473 ± 365 | 15 | 193 ± 235 | 0.22585 | ns |
| **SPA recurrence freq (Hz)** | 0.38 ± 0.34 | 0.36 ± 0.45 | 0.14 ± 0.05 | 0.76 ± 0.6 | 0.05 | 0.39 ± 0.4 | 0.32378 | ns |
| **ISI mean** | 4.7 ± 3.89 | 7.66 ± 5.8 | 7.1 ± 2.38 | 2.22 ± 1.82 | 18.5 | 5.68 ± 4.72 | 0.17386 | ns |
| **LFPg amplitude** | 94.1 ± 92.1 | 181 ± 234 | 64 ± 46.2 | 233 ± 270 | 41.4 | 122 ± 152 | 0.90508 | ns |
| **CSD amplitude** | 79 ± 67.5 | 153 ± 187 | 69.6 ± 37.7 | 189 ± 213 | 34.6 | 104 ± 118 | 0.96788 | ns |
| **MUA amplitude** | 4.59 ± 5.63 | 12.4 ± 9.66 | 13.9 ± 8.63 | 7.65 ± 8.24 | 4.12 | 7.73 ± 7.59 | 0.11288 | ns |

**Supplementary Table 4. Electrophysiological properties of individually classified cell types.** Mean ± SD and sample size (*N*) for 7 electrophysiological properties (metrics) stratified by fine cell-type classification: regularly spiking principal cells (RS-PC,), intrinsically bursting principal cells (IB-PC), and interneurons (IN). Pooled columns for all principal cells (RS-PC + IB-PC combined) and all recorded units are provided for reference. The last 4 columns represent statistical comparisons of electrophysiological metrics across fine cell-type classifications (RS-PC, IB-PC, IN). ns: non-significant, *: p<0.05; **: p<0.01; ***: p<0.001, KW: Kruskal-Wallis ANOVA, MW U: Mann-Whitney U test for all statistics and all tables. Greeen shading shows data for all cells (RS-PC + IB-PC + IN) and all principal cells (RS-PC + IB-PC) pooled. Orange shading marks significance.

| **Metric** | **All cells** | **Principal cells** | **RS-PC** | **IB-PC** | **IN** | **KW_p** | **RS-PC vs IB-PC** | **RS-PC vs IN** | **IB-PC vs IN** |
| --- | --- | --- | --- | --- | --- | --- | --- | --- | --- |
| **N clusters** | **151** | **115** | **99** | **16** | **36** |  |  |  |  |
|  | **Mean ± SD** | **Mean ± SD** | **Mean ± SD** | **Mean ± SD** | **Mean ± SD** |  |  |  |  |
| **Firing Rate (Hz)** | 1.55 ± 1.92 | 1.58 ± 1.99 | 1.66 ± 2.11 | 1.11 ± 0.713 | 1.47 ± 1.73 | 0.87668 | 1.0000 ns | 1.0000 ns | 1.0000 ns |
| **Max Firing Rate (Hz)** | 4.03 ± 4.45 | 3.95 ± 4.62 | 4.12 ± 4.92 | 2.86 ± 1.61 | 4.29 ± 3.93 | 0.83273 | 1.0000 ns | 1.0000 ns | 1.0000 ns |
| **Burstiness (%)** | 0.69 ± 1.85 | 0.83 ± 2.05 | 0.37 ± 1.08 | 3.72 ± 3.74 | 0.23 ± 0.84 | 0.00000 | 0.0000 *** | 0.3264 ns | 0.0000 *** |
| **Burstiness ISI<20 ms (%)** | 16.5 ± 18.4 | 16.2 ± 15.7 | 13.1 ± 13.6 | 35.4 ± 14.1 | 17.5 ± 25.4 | 0.00002 | 0.0000 *** | 0.7574 ns | 0.0063 ** |
| **Half-width 1 (ms)** | 0.25 ± 0.07 | 0.27 ± 0.06 | 0.27 ± 0.06 | 0.3 ± 0.05 | 0.17 ± 0.03 | 0.00000 | 0.1473 ns | 0.0000 *** | 0.0000 *** |
| **Half-width 2 (ms)** | 0.53 ± 0.21 | 0.58 ± 0.18 | 0.57 ± 0.19 | 0.63 ± 0.11 | 0.35 ± 0.18 | 0.00000 | 0.6355 ns | 0.0000 *** | 0.0001 *** |
| **ISI CV** | 1.85 ± 1.26 | 1.77 ± 1.12 | 1.79 ± 1.16 | 1.68 ± 0.9 | 2.12 ± 1.62 | 0.82710 | 1.0000 ns | 1.0000 ns | 1.0000 ns |

**Supplementary Table 5.** Significant changes were observed for several metrics over the different days (D) of the culturing period. D>0=D7, D15, D22 combined. Green shading shows all days pooled. The yellow shading in the bottom table marks significant differences.

| **Metric** | **D0** | **D>0** | **D7** | **D15** | **D22** | **All days** |
| --- | --- | --- | --- | --- | --- | --- |
| **N clusters** | **73** | **77** | **23** | **26** | **28** | **150** |
|  | **Mean ± SD** | **Mean ± SD** | **Mean ± SD** | **Mean ± SD** | **Mean ± SD** | **Mean ± SD** |
| **Firing Rate (Hz)** | 1.71 ± 2.03 | 1.43 ± 1.82 | 2.11 ± 2.26 | 0.6 ± 0.72 | 1.64 ± 1.89 | 1.56 ± 1.93 |
| **Max Firing Rate 10 s (Hz)** | 4.06 ± 4.65 | 4.05 ± 4.3 | 6.1 ± 6.14 | 2.97 ± 2.99 | 3.36 ± 2.82 | 4.05 ± 4.46 |
| **Burstiness (%)** | 0.305 ± 0.798 | 1.08 ± 2.42 | 0.19 ± 0.62 | 1 ± 1.87 | 1.87 ± 3.4 | 0.7 ± 1.85 |
| **Burstiness ISI<20 ms (%)** | 11.5 ± 12.9 | 21.1 ± 21.4 | 8.86 ± 15.7 | 32.2 ± 24.2 | 20.8 ± 17.3 | 16.4 ± 18.4 |
| **Half-width 1 (ms)** | 0.276 ± 0.0816 | 0.23± 0.05 | 0.253 ± 0.06 | 0.21 ± 0.04 | 0.24 ± 0.05 | 0.25 ± 0.07 |
| **Half-width 2 (ms)** | 0.545 ± 0.209 | 0.52 ± 0.2 | 0.58 ± 0.26 | 0.502 ± 0.214 | 0.48 ± 0.14 | 0.53 ± 0.2 |
| **ISI CV** | 1.46 ± 0.756 | 2.22 ± 1.52 | 2.59 ± 1.99 | 2.7 ± 1.32 | 1.47 ± 0.85 | 1.85 ± 1.27 |

Significances for Supplementary Table 5.

| **Metric** | **ANOVA_p** | **KW_p** | **Sig** | **D0 vs D7** | **D0 vs D15** | **D0 vs D22** | **D7 vs D15** | **D7 vs D22** | **D15 vs D22** |
| --- | --- | --- | --- | --- | --- | --- | --- | --- | --- |
| Firing Rate (Hz) | 0.03000 | 0.00051 | *** | 1.0000 ns | 0.0005 *** | 1.0000 ns | 0.0044 ** | 1.0000 ns | 0.0201 * |
| Max Firing Rate (Hz) | 0.06941 | 0.13627 | ns | 0.7088 ns | 1.0000 ns | 1.0000 ns | 0.1648 ns | 0.7364 ns | 1.0000 ns |
| Burstiness (%) | 0.00053 | 0.06276 | ns | 1.0000 ns | 0.5622 ns | 0.6317 ns | 0.2422 ns | 0.2341 ns | 1.0000 ns |
| Burstiness ISI<20 ms (%) | 0.00000 | 0.00001 | *** | 0.1242 ns | 0.0004 *** | 0.1378 ns | 0.0016 ** | 0.0359 * | 0.5667 ns |
| Half-width 1 (ms) | 0.00139 | 0.00226 | ** | 1.0000 ns | 0.0024 ** | 0.3325 ns | 0.1530 ns | 1.0000 ns | 0.4913 ns |
| Half-width 2 (ms) | 0.35048 | 0.21800 | ns | 1.0000 ns | 1.0000 ns | 1.0000 ns | 0.9122 ns | 0.3005 ns | 1.0000 ns |
| ISI CV | 0.00000 | 0.00000 | *** | 0.0825 ns | 0.0000 *** | 1.0000 ns | 1.0000 ns | 0.2511 ns | 0.0001 *** |

**Supplementary Table 6** . Electrophysiological properties in acute (D0) versus cultured slice (D>0) for the previously defined celltypes (RS-PC, IB-PC, IN). Statistically significant cell type differences are highlighted in yellow, all comparisons were carried out with the Mann-Whitney U-test (MW U). Significant values are marked with yellow shading.

| **Metric** | **All cells pooled** | | | **Principal cell** | | | **RS-PC** | | | **IB-PC** | | | **IN** | | |
| --- | --- | --- | --- | --- | --- | --- | --- | --- | --- | --- | --- | --- | --- | --- | --- |
| **N clusters** | 73 | 77 |  | 57 | 58 |  | 50 | 49 |  | 7 | 9 |  | 16 | 19 |  |
|  | **D0 Mean±SD** | **D>0 Mean±SD** | **MW U p** | **D0 Mean±SD** | **D>0 Mean±SD** | **MW U p** | **D0 Mean±SD** | **D>0 Mean±SD** | **MW U p** | **D0 Mean±SD** | **D>0 Mean±SD** | **MW U p** | **D0 Mean±SD** | **D>0 Mean±SD** | **MW U p** |
| **Firing Rate (Hz)** | 1.71 ± 2.03 | 1.43 ± 1.82 | 0.104 ns | 1.72 ± 2.04 | 1.44 ± 1.94 | 0.107 ns | 1.82 ± 2.14 | 1.48 ± 2.1 | 0.033 * | 0.9 ± 0.979 | 1.23 ± 0.444 | 0.091 ns | 1.66 ± 2.05 | 1.39 ± 1.46 | 0.679 ns |
| **Max Firing Rate (Hz)** | 4.06 ± 4.65 | 4.05 ± 4.3 | 0.934 ns | 4.05 ± 4.78 | 3.84 ± 4.5 | 0.589 ns | 4.22 ± 5.01 | 4.02 ± 4.87 | 0.482 ns | 2.81 ± 2.35 | 2.9 ± 0.847 | 0.200 ns | 4.08 ± 4.31 | 4.67 ± 3.69 | 0.508 ns |
| **Burstiness (%)** | 0.3 ± 0.79 | 1.08 ± 2.42 | 0.264 ns | 0.39 ± 0.88 | 1.28 ± 2.69 | 0.514 ns | 0.24 ± 0.62 | 0.5 ± 1.39 | 0.509 ns | 1.46 ± 1.61 | 5.48 ± 4.03 | 0.020 * | 0 ± 0 | 0.453 ± 1.14 | 0.059 ns |
| **Burstiness ISI<20 ms** | 11.5 ± 12.9 | 21.1 ± 21.4 | 0.044 * | 13.8 ± 13.6 | 18.6 ± 17.3 | 0.346 ns | 10.5 ± 9.12 | 15.9 ± 16.7 | 0.559 ns | 38.1 ± 16.2 | 33.3 ± 12.7 | 0.758 ns | 3.02 ± 4.17 | 28.8 ± 30.1 | 0.088 ns |
| **Half-width 1 (ms)** | 0.27 ± 0.08 | 0.23 ± 0.05 | 0.002 ** | 0.3 ± 0.06 | 0.25 ± 0.05 | 0.000 *** | 0.3 ± 0.06 | 0.24 ± 0.04 | 0.000 *** | 0.3 ± 0.06 | 0.29 ± 0.04 | 0.758 ns | 0.17 ± 0.04 | 0.178 ± 0.0245 | 0.908 ns |
| **Half-width 2 (ms)** | 0.54 ± 0.2 | 0.52 ± 0.2 | 0.583 ns | 0.59 ± 0.18 | 0.57 ± 0.18 | 0.882 ns | 0.58 ± 0.18 | 0.57 ± 0.20 | 0.903 ns | 0.67 ± 0.15 | 0.605 ± 0.0663 | 0.210 ns | 0.376 ± 0.2 | 0.345 ± 0.176 | 0.585 ns |
| **ISI CV** | 1.46 ± 0.75 | 2.22 ± 1.52 | 0.000 *** | 1.5 ± 0.79 | 2.03 ± 1.33 | 0.017 * | 1.44 ± 0.77 | 2.13 ± 1.37 | 0.003 ** | 1.92 ± 0.828 | 1.49 ± 0.969 | 0.174 ns | 1.3 ± 0.604 | 2.79 ± 1.93 | 0.003 ** |

**Supplementary Table 7.** Proportion of neurons classified as SPA-responsive across cell types, patients, and days in vitro. A significant chi-squared result indicates that the proportion of event-responsive cells differs significantly across levels of the grouping factor. Significant values are marked with yellow shading.

| **Group** | **Category** | **N** | **‘increased’** | **Ratio of ‘increased’** | **Chi2** | **p_value** | **Significance** |
| --- | --- | --- | --- | --- | --- | --- | --- |
| RS-PC | Cell type (fine) | 99 | 66 | 66.67% | 9.546 | 0.00846 | ** |
| IB-PC | Cell type (fine) | 16 | 15 | 93.75% | 9.546 | 0.00846 | ** |
| IN | Cell type (fine) | 36 | 18 | 50.0% | 9.546 | 0.00846 | ** |
| Principal cell | Cell type (coarse) | 115 | 81 | 70.43% | 4.206 | 0.04028 | * |
| Interneuron | Cell type (coarse) | 36 | 18 | 50.0% | 4.206 | 0.04028 | * |
| HOC26 | Patient | 8 | 4 | 50.0% | 7.015 | 0.07142 | ns |
| HOC27 | Patient | 44 | 35 | 79.55% | 7.015 | 0.07142 | ns |
| HOC29 | Patient | 70 | 40 | 57.14% | 7.015 | 0.07142 | ns |
| HOC30 | Patient | 29 | 20 | 68.97% | 7.015 | 0.07142 | ns |
| 0 | DIV (excl. D30) | 73 | 52 | 71.23% | 8.553 | 0.03586 | * |
| 7 | DIV (excl. D30) | 23 | 9 | 39.13% | 8.553 | 0.03586 | * |
| 15 | DIV (excl. D30) | 26 | 17 | 65.38% | 8.553 | 0.03586 | * |
| 20 | DIV (excl. D30) | 28 | 20 | 71.43% | 8.553 | 0.03586 | * |

**Pairwise Spearman correlations between electrophysiological metrics**

Correlation matrices for all cells pooled and for each cell-type subgroup are provided in Suppl. Figure 1. Across all 151 units, firing rate and maximum firing rate were strongly positively correlated (ρ = 0.868, p < 0.001), as expected, given that both reflect overall discharge intensity. Burstiness % and burstiness ISI<20 ms were moderately correlated (ρ = 0.551, p < 0.001), confirming partial but non-redundant measurement of burst propensity by the two indices. Burstiness ISI<20 ms also assesses the interneurons with high firing rate, i.e. the fast-spiking cells. ISI CV was significantly associated with maximum firing rate (ρ = 0.331, p < 0.001) and burstiness ISI < 20 ms (ρ = 0.458, p < 0.001), consistent with high ISI variability being driven partly by burst–pause alternation. The two half-width metrics were substantially correlated with each other (ρ = 0.447, p < 0.001) and were largely independent of rate and burstiness metrics, supporting their use as orthogonal waveform-shape features.

Within RS-PCs (n = 99), the correlation structure was qualitatively similar to the pooled dataset but with stronger burstiness associations. Within IB-PCs (n = 16), the pattern diverged substantially: firing rate correlated strongly with maximum firing rate (ρ = 0.719, p = 0.002) but showed negative associations with half-width 1 (ρ = −0.335) and burstiness ISI < 20 ms correlated negatively with half-width 1 (ρ = −0.529, p = 0.035) and positively with ISI CV (ρ = 0.600, p = 0.014), suggesting that among IB-PCs, higher burst propensity was associated with narrower, more irregular-firing cells. Within INs (n = 36), the dominant relationships were between firing rate and maximum firing rate (ρ = 0.889, p < 0.001) and between burstiness ISI < 20 ms and ISI CV (ρ = 0.586, p < 0.001), while waveform metrics were uncorrelated with rate or burst metrics, consistent with the electrophysiological homogeneity of the interneuron population at the half-width level.

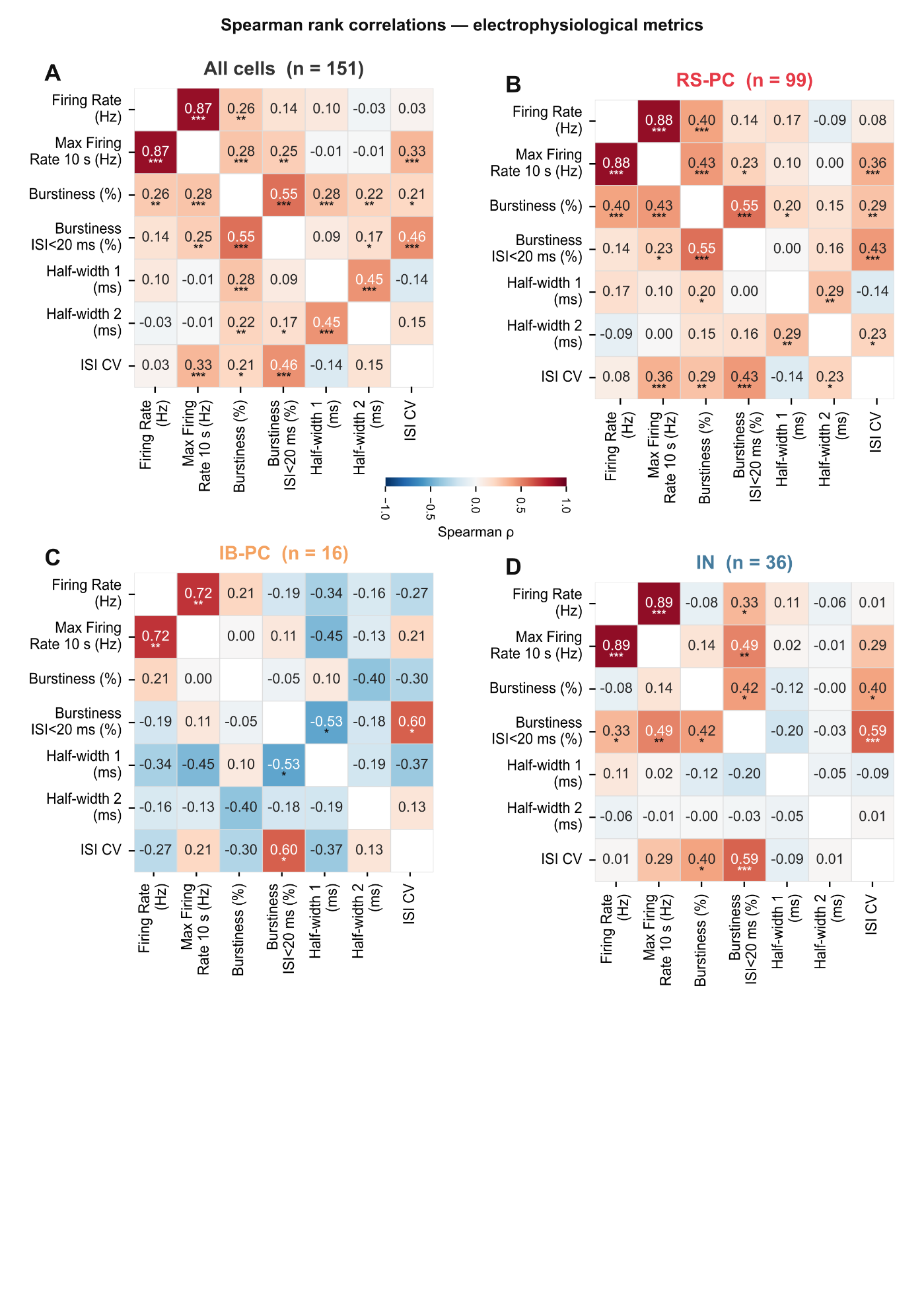
**Supplementary Figure 1. Correlations between electrophysiological properties of the different clustered cell types.**

A) Correlations for all cells combined, B) for RS-PCs, C) for IB-PCs and D) for INs. Note the different patterns for IB-PCs than for the other cell types. For more explanation see the text.

**Wide-field imaging**

Seventeen organotypic slices were recorded with wide-field imaging. Fourteen slices were injected with viruses to induce expression of the Ca^2+^-sensor GCaMP. One slice was in a well together with a virus injected slice, and we observed GCaMP-positive cells in this slice, mediated by the culture medium. Sixteen slices were recorded multiple times, at different time points during the culturing period (see Suppl. Table 8). We saw GCaMP-related Ca^2+^-signals in all slices with viral injection. We analysed the calcium signal for each slice, all over their culturing period. Fluorescence originating from GCaMP+ cells was usually increased with time (in n=10/13 slices examined at several time points), both in intensity and in the spatial dimension, invading larger areas of the slice with time (see Suppl. Figure 2). Slices with low virus concentration were not expressing this spatial and intensity increase (n=2 slices with 1:10, n=1 slice with 1:100 and the neighbouring slice without injection).

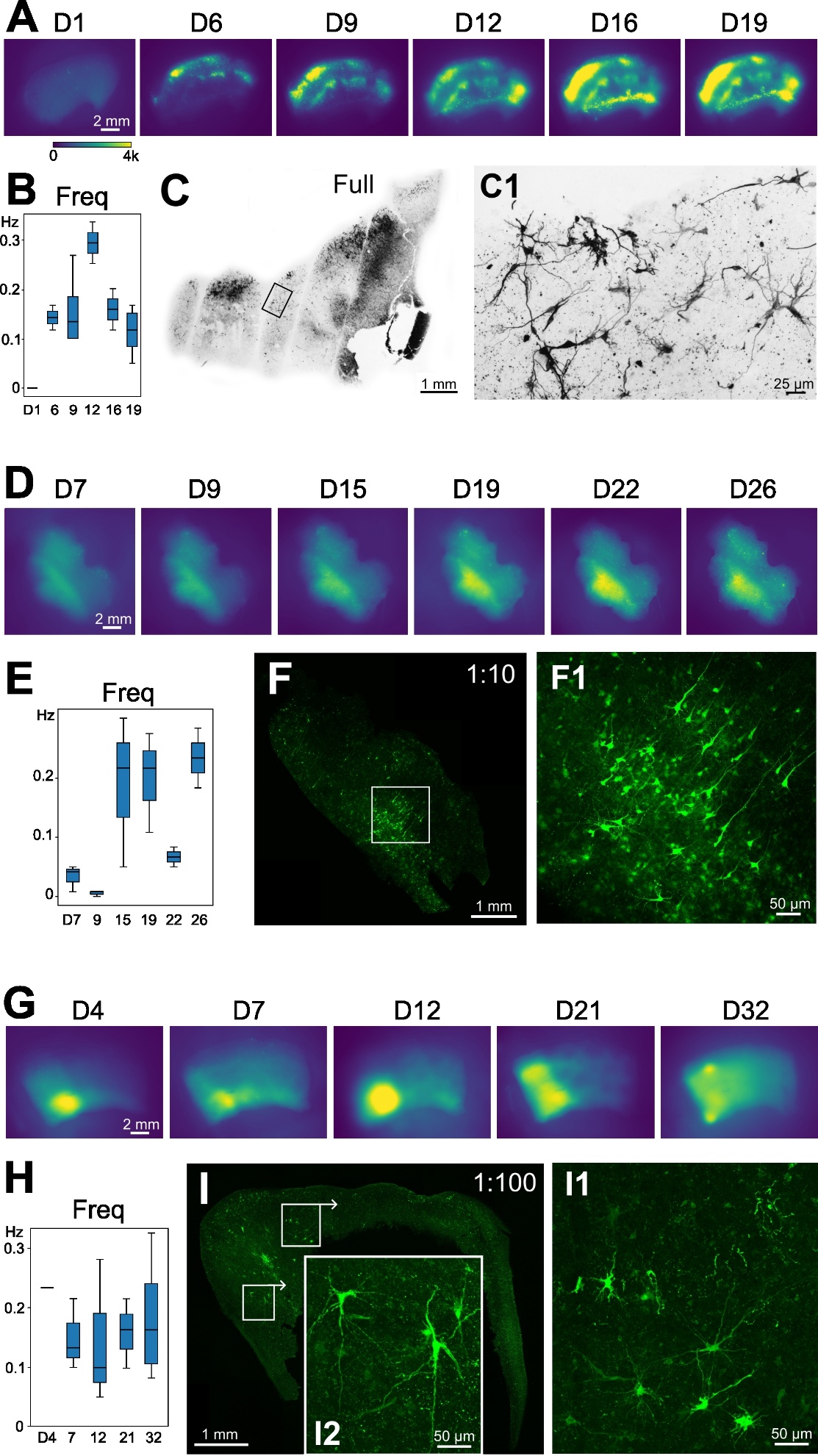

**Supplementary Figure 2. Wide-field imaging of human organotypic slices.**

**(A)** Time-averaged mean fluorescence images of slice HOC13w2, injected with undiluted virus of EF1a-GCaMP6s-WPRE (PHP.eBb, titer: 2.47E+13 vg/ml). GCaMP expression gradually increased from D6 to D19 both in intensity and spatially, invading larger areas with time. No expression was observed at D1. Warm colours (green-yellow) indicates higher GCaMP expression (0-4000, arbitrary unit, the same colour scale applies to all images on this figure). **(B)** Calcium transient recurrence frequency (Freq) for the slice shown on (A). Note that this value is 0 at D1. **(C)** Immunostaining against GCaMP with a peroxidase-based method of the same slice. The boxed region is shown with higher magnification (C1). GCaMP signal appears as dark labelling of neurons with long dendrites. The slice was fixed on D21. **(D)** Time-averaged mean fluorescence images of slice HOC15p2w3, injected with 1:10 virus dilution of EF1a-GCaMP6s-WPRE (PHP.eBb, original titer: 2.47E+13 vg/ml). The increase in GCaMP expression is not as considerable, as in the slice on (A). **(E)** Frequency of the calcium transients emerging in slice shown on (D) is low at the beginning of the culturing period (D7-D9), then increases and stays high until the end. **(F)** Native fluorescence image of a section from the slice sjown on (D), fixed on D35. The boxed region is magnified in F1. GCaMP-positive neurons with pyramidal morphology are visible. **(G)** Time-averaged mean fluorescence images of slice HOC21p2w3s2, injected with a 1:100 virus dilution of Syn-jGCaMP7s-WPRE (PHhP.eBb, original titer: 2.82E+14 vg/ml). GCaMP expression is shown over the culturing period. **(H)** The frequency of the populational calcium transients was stable all over the culturing period. **(I)** Native fluorescence image of the slice shown on (G). The area outlined with the larger box is magnified on I1, the smaller on I2. Note the large neurons with long dendrites on both magnified panels.

**Supplementary Table 8. Wide-field imaging of organotypic slices at different time points after viral injection time (D0 to D31).** Frequency of calcium transients (Hz, mean ± st.dev.) measured over the entire slice are shown for each slice at each recording day. Empty cells denotes no recording from that slice on that day. One number means that the frequency of the calcium transients was determined from the only video made from the given slice on the given day, mean ± st.dev. denotes that frequency was calculated from multiple recordings of the given slice on the same day.

| **Sample** | **Slice** | **Virus cc** | **D1** | **D4** | **D6** | **D7** | **D9** | **D12** | **D15** | **D16** | **D19** | **D21** | **D22** | **D26** | **D31** |
| --- | --- | --- | --- | --- | --- | --- | --- | --- | --- | --- | --- | --- | --- | --- | --- |
| HOC13  EF1a-GCaMP6s-WPRE  (PHP.eB, titer: 2.47E+13 vg/ml) | w1s1 | no | 0.0 ± 0.0 |  | 0.0 |  |  | 0.0 |  | 0.0 | 0.0 |  |  |  |  |
|  | w2 | full | 0.0 ± 0.0 |  | 0.142 ± 0.036 |  | 0.157 ± 0.071 | 0.293 ± 0.059 |  | 0.159 ± 0.059 | 0.114 ±0.048 |  |  |  |  |
|  | w3 | full | 0.0 ± 0.0 |  | 0.064 ± 0.051 |  | 0.067 ± 0.017 | 0.207 ± 0.073 |  |  |  |  |  |  |  |
|  | w5s1 | full | 0.0 ± 0.0 |  | 0.184 |  | 0.075 ± 0.056 | 0.017 |  |  |  |  |  |  |  |
|  | w5s2 | full on neighboring slice | 0.0 ± 0.0 |  |  |  | 0.017 | 0.0 |  | 0.112 ± 0 .059 | 0.017 |  |  |  |  |
|  | w6 | full | 0.0 ± 0.0 |  | 0.067 ± 0.047 |  |  |  |  |  |  |  |  |  |  |
| HOC15  EF1a-GCaMP6s-WPRE  (PHP.eB, titer: 2.47E+13 vg/ml) | p1w1 | 1:10 |  |  |  | 0.022 ± 0.026 | 0.029 ± 0.018 |  | 0.028 ± 0.035 |  | 0.042 ± 0.035 |  | 0.050 ± 0.047 | 0.050 ± 0.017 |  |
|  | p1w3 | full |  |  |  | 0.053 ± 0.078 | 0.025 ± 0.020 |  | 0.065 ± 0.043 |  | 0.086 ± 0.070 |  | 0.194 ± 0.100 | 0.368 ± 0.047 |  |
|  | p2w1 | full |  |  |  | 0.008 ± 0.0 | 0.028 ± 0.017 |  | 0.059 ± 0.059 |  | 0.017 |  | 0.033 ± 0.000 | 0.042 ± 0.012 |  |
|  | p2w2 | no |  |  |  | 0.0 ± 0.0 | 0.0 ± 0.0 |  | 0.0 ± 0.0 |  | 0.0 ± 0.0 |  | 0.0 ± 0.0 | 0.0 ± 0.0 |  |
|  | p2w3 | 1:10 |  |  |  | 0.033 ±0.022 | 0.006 ± 0.005 |  | 0.190 ± 0.128 |  | 0.200 ± 0.085 |  | 0.067 ± 0.017 | 0.234 ± 0.050 |  |
| HOC21  Syn-jGCaMP7s-WPRE  (PHP.eB, titer: 2.82E+14 vg/ml) | p2w1s1 | full |  | 0.008 ± 0.012 |  | 0.254 ± 0.085 |  | 0.224 ± 0.012 |  |  |  | 0.386 ± 0.077 |  |  |  |
|  | p2w1s2 | full |  | 0.167 ± 0.058 |  | 0.203 ± 0.147 |  | 0.131 ± 0.132 |  |  |  | 0.214 ± 0.035 |  |  | 0.263 ± 0.212 |
|  | p2w2s1 | 1:10 |  | 0.067 ± 0.024 |  | 0.124 ± 0.012 |  | 0.199 ± 0.060 |  |  |  | 0.216 ± 0.108 |  |  | 0.090 ± 0.038 |
|  | p2w2s2 | 1:10 |  | 0.050 |  | 0.058 ± 0.044 |  | 0.042 ± 0.021 |  |  |  | 0.083 ± 0.047 |  |  |  |
|  | p2w3s1 | 1:100 |  | 0.050 |  | 0.044 ± 0.010 |  | 0.077 ± 0.048 |  |  |  | 0.060 ± 0.050 |  |  |  |
|  | p2w3s2 | 1:100 |  | 0.234 |  | 0.149 ± 0.060 |  | 0.144 ± 0.122 |  |  |  | 0.158 ± 0.059 |  |  | 0.189 ± 0.102 |
| **Total N of slices** | |  | **6** | **6** | **5** | **11** | **9** | **11** | **5** | **3** | **8** | **6** | **5** | **5** | **3** |

**Histology**

**NeuN**

The variability of the neuronal morphology and survival was high in the different organotypic cultures. We found very different neuronal morphology and densities in the different samples, fixed on the same day but originating from different patients (Suppl. Fig 3A). For example, at D21, numerous neurons survived and the layers of the neocortex could be easily distinguished in certain samples, but very few neurons were preserved and the layer borders faded away in other samples.

We found a large variability in the neuron numbers linked to the location of the section within the depth of the slice. Usually, sections originated from the surface of the slice had less cells, than sections in the middle of the slice (Suppl. Fig. 3).

*Day-by-day vs. week-by-week changes*

Organotypic cultures were fixed at numerous different days during the culturing period ranging from D0 to D42. Day-by-day analysis of the NeuN density means when all different days were considered as different time point and compared to each other with Kruskall-Wallis ANOVA. In the week-by-week changes we made groups including data derived from the end of each week, such as D7±1, D14±1. D21±1, D28±1 and D35±1 and compared to each other with ANOVA.

*Viral treatment*

Viral treatment usually affected both the neuronal density and the morphology. The site of the injection could be frequently outlined in the NeuN-stained section: a dramatic neuron loss and tissue damage was observed in a large area around the injection site. In some cases we found NeuN-stained cells with pathological morphology, and signs of cell degradation were also observed (Suppl. Fig. 3B). Cells were usually more preserved, and had healthier-looking morphology at distant sites from the injection.

*hCSF vs. AM*

Slices derived from 11 samples were cultured in human cerebrospinal fluid (hCSF), slices from 16 samples were treated with artificial medium (AM), whereas in one sample we tested both media (see Suppl. Table 2). The use of AM gave better results in tissue and cell preservation. The number of NeuN+ cells showed a stronger decrease with time in slices treated with hCSF than with AM, although the differences were not significant (p>0.05, Kruskal-Wallis ANOVA).

*Antibiotics/Antimicotics treatment*

We tried different combinations of antibiotic and antimycotic treatments when using hCSF medium. Antibiotics penicillin and streptomycin (PS) were used for the first week of culture period, and this treatment was supplemented with the fungicid amphotericin B (PSA) in 4 out of 11 samples. Slices treated with PSA showed higher numbers NeuN+ cells, than slices treated only with PS (p<0.01, Two-tailed unpaired t-test), already after 2-3 days of culturing.

In n=3 samples we also tried to maintain the human organotypic cultures without PS/PSA treatment. These slices had lower neuronal density and had neurons with pathological morphology and cellular debris (Suppl. Fig. 3D), than their sister slices treated with PSA. Furthermore, these slices often showed signs of infection after a short to medium (D3-D21) culture time.

*Epi vs. tumor*

Densities of NeuN+ cells derived from epileptic patients were compared to those obtained from non-epileptic tumour patients. When comparing all slices (from D0 to D42), no differences were observed between slices derived from epileptic or non-epileptic tissues (unpaired t-test, n(epi)=8, n(non-epi)=7 patients, p=0.2381). Next, we also checked whether differences existed in the acute samples (only at D0), and found that neuronal density was similar (unpaired t-test, n(epi)=5, n(non-epi)=8 patients, p=0.1506).

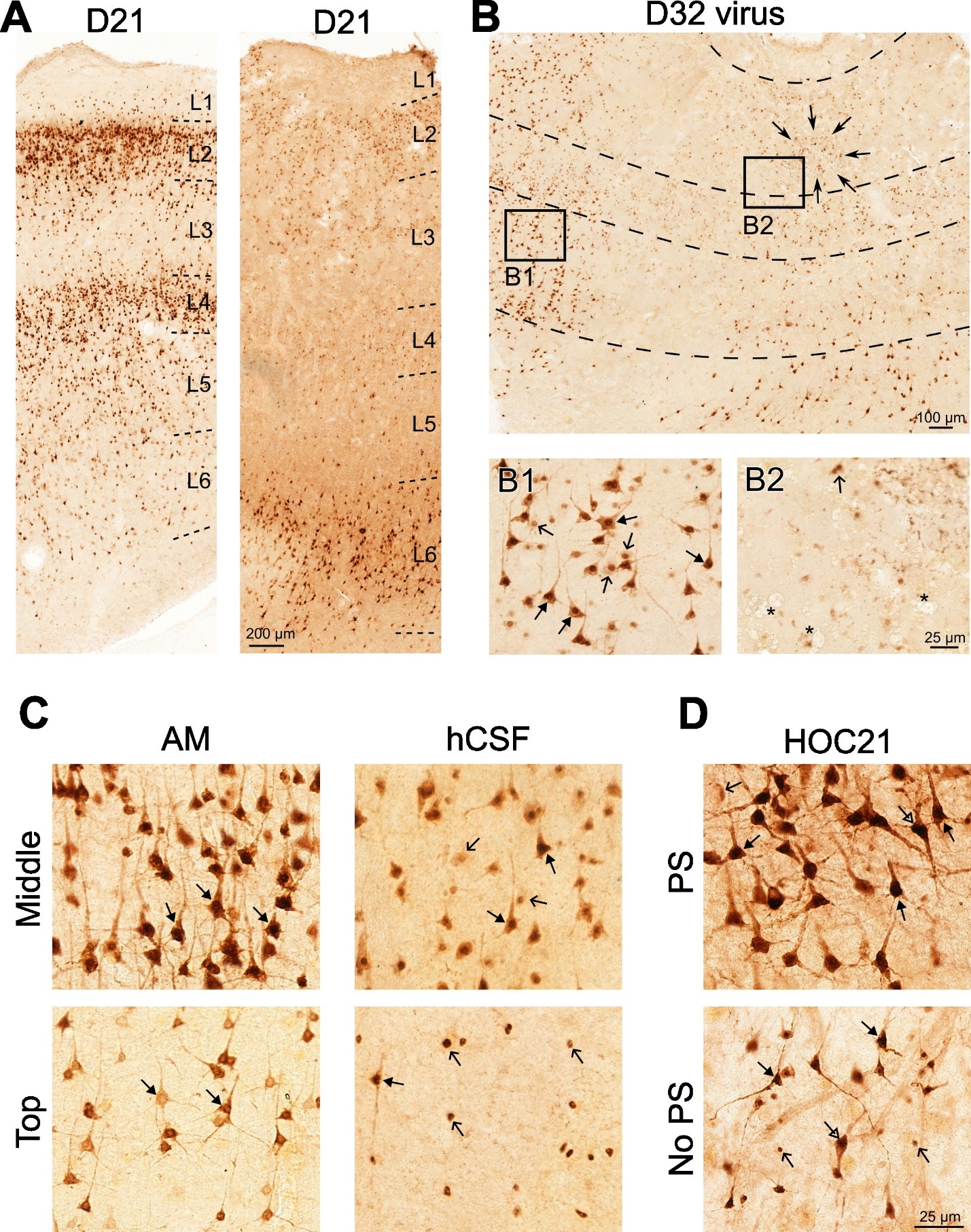

**Supplementary Figure 3. NeuN staining**

**(A)** Neuronal density shows a substantial variability between samples cultured for the same period. At D21 numerous neurons are preserved in the tissue sample HOC25 (left), whereas most neurons are lost at D21 in the sample HOC24 (right). The layers can be distinguished in both cases, but very few and only lightly stained neurons are present in L1-5 of the HOC24 culture.

**(B)** This image shows a NeuN-stained section of a virally injected slice fixed at D32. The injection site is outlined by arrows. Scattered neurons are visible around the injection site (enlarged in B2), whereas more neurons are preserved at distant sites (B1). Numerous neurons with pathological morphology (open arrows) are present among the healthy-looking neurons (arrows). Note the holes in the tissue at the injection site (asterisks on B2).

**(C)** The effect of the culturing medium, as well as of the position of the section within the slice is visible both at the level of the neuronal density and the morphology. These images derive from two D2 organotypic cultures, one kept in AM (left side), the other in hCSF (right side). Considerably higher number of neurons survived in AM than in hCSF, and they all look healthy (arrows). In contrast, in the slice kept in hCSF, the neuronal density is lower and beside the healthy-looking neurons (arrows) cells with pathological or degrading morphology (open arrows) are present. Sections originating from the middle of the slice (upper images) had higher neuronal density than those derived from the surface (bottom images). Neurons look healthier in AM than in hCSF, even at the surface of the slice. Several very small NeuN-stained nuclei (open arrows) were present on the surface of the slice kept in hCSF.

**(D)** This image shows the difference in the NeuN staining when Penicillin/Streptamycin treatment was used (PS) compared to slices without any antibacterial treatment (No PS). Large number of healthy-looking neurons (arrows) are present in the slice treated with PS (fixed at D32), whereas lower neuron density and more neuronal debris/degrading neurons (open arrows) can be observed in the slice without treatment (fixed at D29). Note, that few neurons with upside-down orientation (open triangle arrows on both images) are visible at this late stage of culturing period.

**GFAP**

To see the inhomogeneity of the glial coverage within the slice, we examined the standard deviation (SD) of the glial coverage in the different sections originated from a given slice (n=63 cultured slices, n=5 acute slices). Over the culturing period the mean SD value increased linearly 2-fold (D0-37) in general. The different culturing conditions did not cause significant changes in the course of the gliosis: n(hCSF)=21, n(AM)=10, p=0.9834; neither the viral treatment compared to control slices: n(ctrl)=23, n(v)=18, p=0.2348.

Although the viral treatment did not have significant effect on the glial coverage, the mean value of the control cases was 2/3 of the viral treated. Same applies to the analysis of the different culturing media: the difference was not significant between AM and hCSF, but the mean value of the AM was 2-fold compared to the hCSF.

This indicates that the lowest gliosis was observed in slices kept in AM and treated with complex antibiotic and antifungal treatment. Viral injection induced a more severe gliosis compared to non-injected slices.

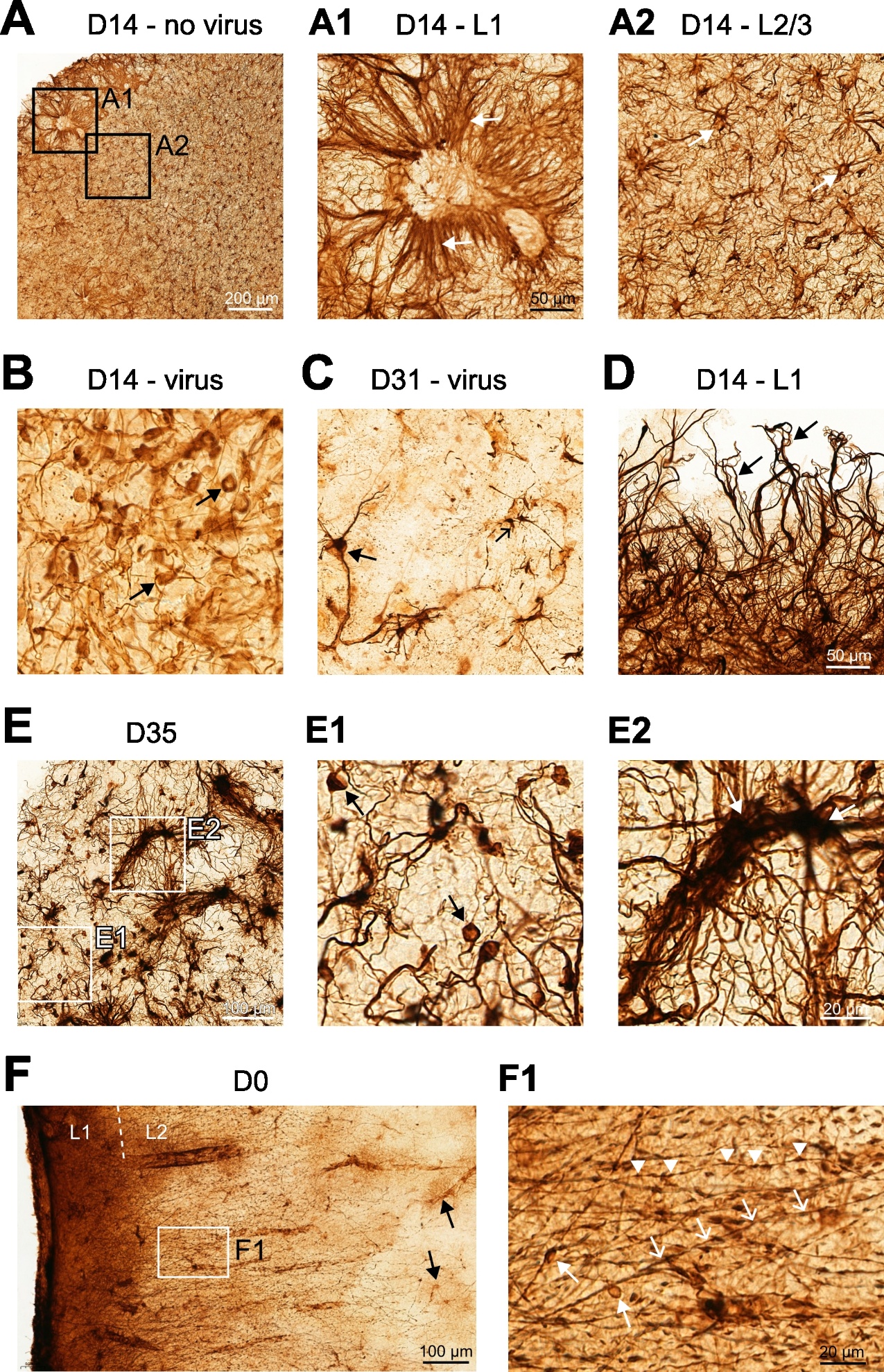

**Supplementary Figure 4. GFAP staining**

A) Very different GFAP+ structures were observed in the different layers of the organotypic neocortical slices. In layer 1 (L1, A1), a dense network of very long astroglial processes (white arrows) was seen, whereas in layers 2/3 (A2, L2/3), the typical activated astroglial cells (white arrows) formed a gliotic network.

B) In slices with viral injection, astroglial cells with unusual morphology were observed. In this D14 culture round cell bodies with no or with only a few, moderately stained processes (black arrows) were present all over the slice.

C) In virally transduced slices, after several weeks of culture, the loss of astroglial cells was observed. In this slice fixed at D31, only scattered GFAP+ cells were visible. Some were hypertrophic with long, thick processes (arrow), while others possessed a small cell body and thin processes (open arrow).

D) Reactive astroglial cells located in layer 1 (L1) stretch out their processes out from the slice (fixed on D14).

E) In long-kept organotypic slices (after several weeks of culture), different glial cells and GFAP+ structures were seen. In this D35 slice small, strongly stained cells with only few processes (E1, arrows) were seen in the vicinity of large, strongly stained blood vessel-looking structures (E2, white arrows).

F) In several cases long GFAP+ processes were observed stretching from L1 towards the deeper layers. Only scattered astroglial cells were visible in L2/3 (arrows on the left image), whereas the blood vessels were covered by darkly stained endfeet. The inset is magnified on F1. Long and straight processes were seen (open arrows), and some of them had beaded structures (arrowheads point on these beads). Sometimes larger bead-like structures were also detected (arrows).

**IBA1**

The IBA1 coverage of the slices derived from the different patients was similar when we compared slices without viral treatment (HOC comparison, Kruskal-Wallis ANOVA, (no viral): ns, p=0.3566, number of HOC: 13, n=37 slices). The microglial activation (the perimeter-to-area ratio, PAR) was also similar in the different HOC samples (HOC comparison (no viral): ns, p=0.2438, number of HOC: 13, n=37 slices). When comparing the slices day-by-day, we could not demonstrate significant differences neither in the coverage (day-by-day changes, Kruskal-Wallis ANOVA, (no viral): ns, p=0.3015, number of days: 20, n=37 slices), nor in the PAR (day-by-day changes (no viral): ns, p=0.0553, number of days: 20, n=37 slices). When comparing the slices at the end of each week (D7±1, D14±1, D21±1, etc.), we found that the IBA1 coverage was similar (Kruskal-Wallis ANOVA, p=0.2641), but the PAR was significantly different (Kruskal-Wallis ANOVA, p=0.0029). The post-hoc test showed that D14±1 was significantly different from D0, p=0.0047).

The coverage of the slices by IBA1-stained cells did not show significant differences over the culture period. However, the standard deviation of the coverage values within the slice began to increase most strikingly after D21 (Suppl. Fig. 5A). The standard deviation of the perimeter-to-area ratio (PAR) within the field of interest shows the differences among the detected cells. Based on this, the size of the somas showed higher variability after D7 (Suppl. Fig. 5B).

When we compared the slices kept in AM to those in hCSF, we found that the mean IBA1-coverage was slightly but not significantly higher in AM than in hCSF (Mann-Whitney test, n(hCSF)=6, n(AM)=8, p=0.06620). The microglial activation was significantly lower in AM than in hCSF, assessed by the PAR (Mann-Whitney test, n(hCSF)=6, n(AM)=8, p=0.0027). This suggests, that a slightly reduced number of microglial cells with higher activation levels are present in hCSF compared to AM.

We verified the effect of viral treatment by comparing four pairs of slices at different time points (from D32 to D42). One slice of the pair was transduced with a virus, whereas the other served as a control. Virus treated slices were similar to non-treated ones in their IBA1-coverage (paired t-test, p=0.9296, Suppl. Fig. 5D), and looked similar at light microscopic level, however, the mean PAR was slightly but not significantly higher (paired t-test, p=0.25), showing a higher activation level of microglial cells linked to viral treatment. This also implies that microglial cells in control slices have smaller cell bodies and more and longer processes, than in viral treated slices.

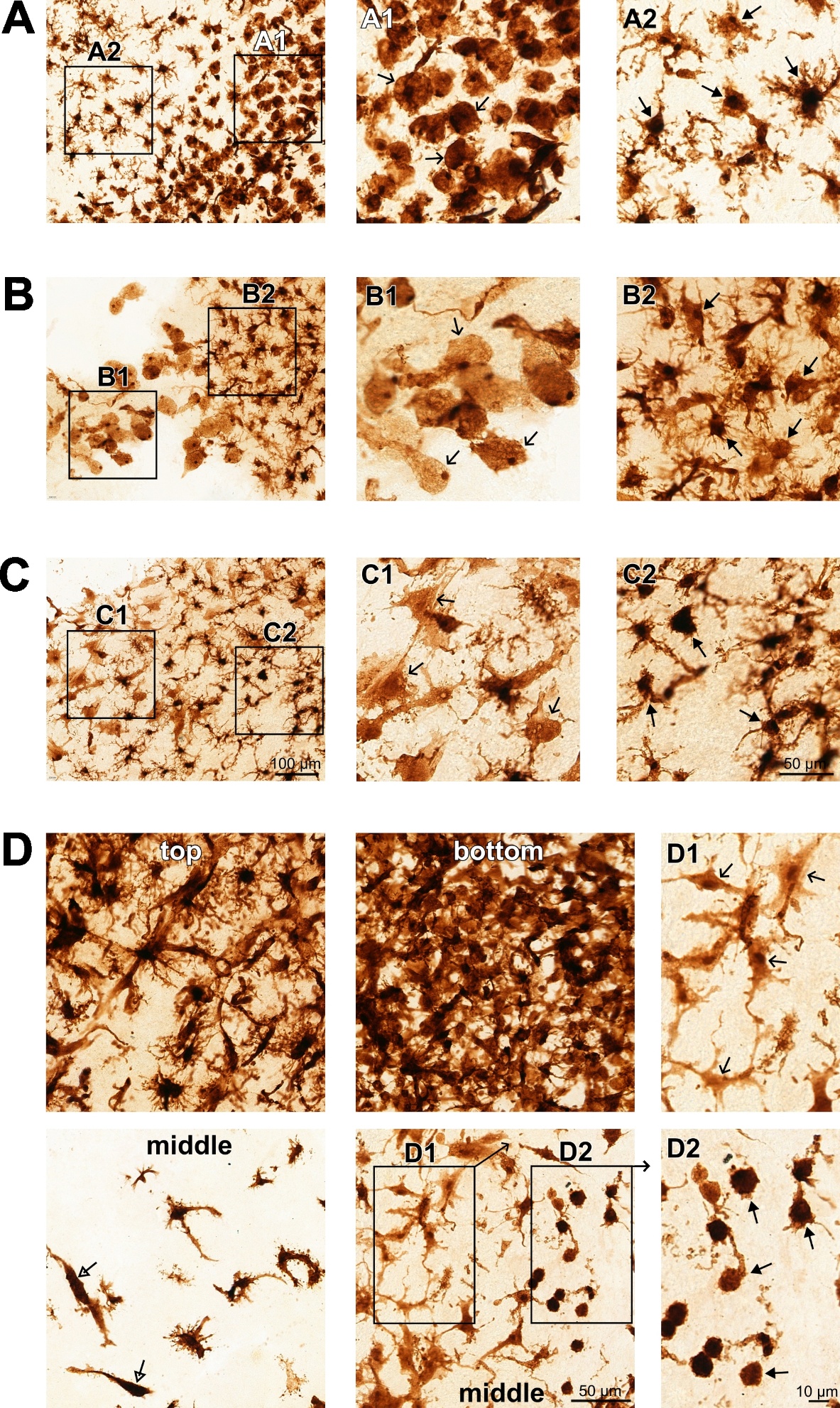

**Supplementary Figure 5. IBA1 staining**

**(A)** Patches of IBA1-positive microglial cells with different morphology in a D21 culture. On the right side (magnified on A1) phagocytic forms of microglial cells are visible (open arrows), whereas on the left side (magnified on A2) reactive microglial cells are present with large cell body and numerous thick processes (arrows).

**(B)** Similar patches of phagocytic (B1, open arrows) and reactive (B2, arrows) microglial cells were observed in a D42 culture. Note, that the amoeboid microglial cells are considerably larger and less heavily stained, than at D21 (same scale).

**(C)** Large patch of microglial cells having an intermediate morphology between reactive and amoeboid types (C1, open arrows), having large, homogeneously lightly stained cell body and several thick and thin processes, from a D21 slice. Next to this patch of cells typical reactive microglial cells are visible (C2, arrows).

**(D)** Considerable differences were seen in the IBA1-coverage of the sections derived from the surfaces (top or bottom, upper row) or the middle (lower row) of the organotypic culture. The left panel shows a slice culture fixed at D32, the middle panel shows a D35 slice. Very dense microglial network was found on the top and the bottom of the slices, whereas a sparse to moderate number of microglial cells were visible in the middle of the slices. Note the rod-like microglial cells in the middle of the D32 slice (open triangle arrows). In the middle of the D35 slice patches of IBA1+ cells with different morphology were present next to each other. The marked areas are magnified on D1 and D2. Lightly stained microglial cells with large cell bodies and long thick and thin processes are shown on D1 (open arrows), and densely stained cells with small and round cell bodies are on D2 (arrows). Some of these cells do not have any processes, others have only few, thin processes.

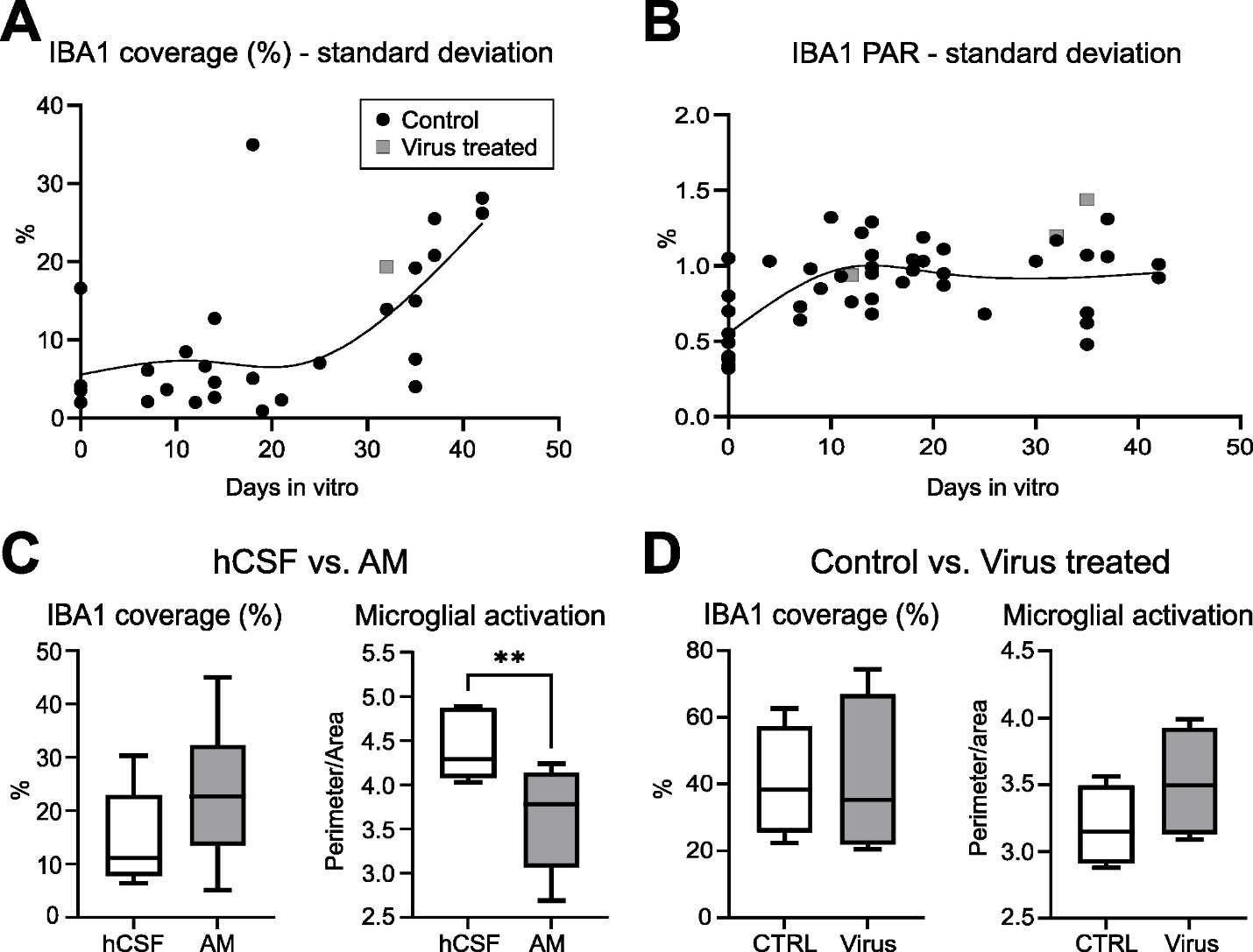

**Supplementary Figure 6. Microglial coverage and activation**

**(A)** Standard deviation of the IBA1 coverage values within the examined slices, during the course of the culture period. Note that after D30 standard deviation considerably increases, pointing to the microglial coverage variability within slices. This might be related to the formation of a microglial sheet on the two surfaces of the slices.

**(B)** The standard deviation of the perimeter-to-area ratio (PAR) within the examined slices notably increased after D7, and stayed high along the culturing period. This suggests that an inhomogeneous distribution and/or a large variability of the different types of microglial cells are present after the first week of culture.

**(C)** The effect of culture medium on the IBA1-coverage (left) and the microglial activation level (assessed by the PAR, right). IBA1-coverage was slightly, but not significantly higher in artificial medium (AM) compared to human cerebrospinal fluid (hCSF), whereas PAR was significantly lower in AM than in hCSF.

**(D)** Viral treatment did not have significant differences neither on the IBA1-coverage of the slices, nor on the microglial activation. However, viral treatment had a tendency of activating more microglial cells, i.e. PAR was slightly but not significantly higher in virus treated slices, compared to non-treated ones.

**Parvalbumin**

Considering the temporal changes across conditions, culturing in hCSF had the greatest impact on cell density (Kruskal-Wallis ANOVA, n=11, *, p=0.0109), samples cultured in AM, whether included the viral-treated ones or not, were not significantly different (Kruskal-Wallis ANOVA, n(ctrl)=27, p1=0.2073, n(virus)=32, p2=0.0830). The tissues cultured in AM without viral injection had the highest mean value, and its lower quartile was higher than zero. In the case of either hCSF or viral-treated samples the mean value was lower, with numerous zero among the values, and their lower quartiles were also zero. The effect of the antimycotic treatment in hCSF was negligible, as the mean and quartiles were similar for PS and PSA.

**Calretinin**

Although based on the data, differences are not significant across the different culturing properties, comparing the mean values, both in the case of viral treatment and culturing media, the control case and the AM resulted in a two-fold density compared to the viral-treated or hCSF-cultured tissues. Furthermore, in the tissues cultured in hCSF, the total disappearance of the CR+ cells was more typical. In the viral-treated tissue, this phenomenon was not typical; in 1 of 5 data pairs, the density was 3-fold higher in the viral-treated tissue than in its control.

Comparing the change in the PV+ and CR+ cell density: although the cell density dropped after two weeks of culturing in both cases, the total disappearance of the CR+ cells was less typical. CR+ staining was more robust, the mean values were reduced to the half of the original values until the 3rd week, then stayed stable. In contrast, the mean value of PV+ cell density was one-third of the control after one week, and after two weeks of culturing total disappearance was more typical.

**Supplementary Table 9. Quantitative morphological analysis.**

D=days in vitro; n= number of slices examined; SD=standard deviation; hCSF: human cerebrospinal fluid; AM: arttificial medium; PS: penicillin + streptomycin; PSA: penicillin + streptomycin + amphotercinB

|  |  |  | **NeuN cell density (cells/mm2)** | | | | | | | | | | | |
| --- | --- | --- | --- | --- | --- | --- | --- | --- | --- | --- | --- | --- | --- | --- |
|  | **Total** | | **hCSF** | | **AM** | | **not treated** | | **viral treated** | | **PS (hCSF)** | | **PSA (hCSF)** | |
| **D** | **n** | **mean±SD** | **n** | **mean±SD** | **n** | **mean±SD** | **n** | **mean±SD** | **n** | **mean±SD** | **n** | **mean±SD** | **n** | **mean±SD** |
| 0 | 17 | 1068 ± 333 | 6 | 1094 ± 345 | 11 | 1054 ± 342 | 15 | 1055 ± 353 | 0 |  | 2 | 1170 ± 76 | 4 | 1056 ± 437 |
| 7±1 | 19 | 617 ± 221 | 11 | 542 ± 208 | 8 | 719 ± 207 | 14 | 685 ± 191 | 1 | 724 | 4 | 349 ± 112 | 7 | 653 ± 162 |
| 14±1 | 23 | 605 ± 255 | 10 | 457 ± 196 | 13 | 718 ± 241 | 12 | 546 ± 243 | 6 | 802 ± 138 | 3 | 398 ± 179 | 7 | 483 ± 211 |
| 21±1 | 13 | 684 ± 347 | 2 | 416 ± 263 | 11 | 733 ± 347 | 6 | 573 ± 151 | 4 | 991 ± 460 | 1 | 230 | 1 | 602 |
| 28±1 | 8 | 608 ± 360 | 0 |  | 8 | 608 ± 360 | 3 | 537 ± 502 | 4 | 707 ± 324 | 0 |  | 0 |  |
| 35±1 | 7 | 712 ± 526 | 0 |  | 7 | 712 ± 526 | 5 | 532 ± 327 | 2 | 1162 ± 816 | 0 |  | 0 |  |
| 37+ | 6 | 516 ± 156 | 0 |  | 6 | 516 ± 156 | 2 | 419 ± 88 | 4 | 564 ± 169 | 0 |  | 0 |  |

|  |  |  | **GFAP coverage (%)** | | | | | | | | | | | |
| --- | --- | --- | --- | --- | --- | --- | --- | --- | --- | --- | --- | --- | --- | --- |
|  | **Total** | | **CSF** | | **AM** | | **not treated** | | **viral treated** | | **PS (hCSF)** | | **PSA (hCSF)** | |
| **D** | **n** | **mean±SD** | **n** | **mean±SD** | **n** | **mean±SD** | **n** | **mean±SD** | **n** | **mean±SD** | **n** | **mean±SD** | **n** | **mean±SD** |
| 0 | 20 | 39.35 ± 17.84 | 9 | 52.64 ± 14.50 | 11 | 28.47 ± 12.14 | 14 | 30.15 ± 11.52 | 0 |  | 6 | 60.81 ± 8.62 | 3 | 36.29 ± 7.32 |
| 7±1 | 16 | 25.50 ± 16.80 | 12 | 21.63 ± 17.32 | 4 | 37.12 ± 8.52 | 10 | 26.21 ± 15.63 | 1 | 11.52 | 5 | 26.90 ± 21.37 | 7 | 17.87 ± 14.34 |
| 14±1 | 25 | 34.74 ± 15.96 | 12 | 30.26 ± 19.95 | 13 | 38.87 ± 10.29 | 10 | 34.51 ± 18.70 | 8 | 36.83 ± 14.68 | 5 | 25.78 ± 11.28 | 7 | 33.46 ± 24.82 |
| 21±1 | 9 | 39.07 ± 21.43 | 1 | 70.39 | 8 | 35.16 ± 19.16 | 5 | 38.25 ± 26.29 | 2 | 25.16 ± 2.32 | 0 |  | 1 | 70.39 |
| 28±1 | 6 | 22.47 ± 6.91 | 0 |  | 6 | 22.47 ± 6.91 | 1 | 29.18 | 5 | 21.13 ± 6.80 | 0 |  | 0 |  |
| 35±1 | 6 | 47.44 ± 10.02 | 0 |  | 6 | 47.44 ± 10.02 | 5 | 49.59 ± 9.54 | 1 | 36.73 | 0 |  | 0 |  |
| 37+ | 6 | 22.29 ± 12.42 | 0 |  | 6 | 22.29 ± 12.42 | 2 | 19.98 ± 23.39 | 4 | 23.45 ± 8.32 | 0 |  | 0 |  |

|  |  |  | **IBA1 coverage (%)** | | | | | | | | | | | |
| --- | --- | --- | --- | --- | --- | --- | --- | --- | --- | --- | --- | --- | --- | --- |
|  | **Total** | | **CSF** | | **AM** | | **not treated** | | **viral treated** | | **PS (hCSF)** | | **PSA (hCSF)** | |
| **D** | **n** | **mean±SD** | **n** | **mean±SD** | **n** | **mean±SD** | **n** | **mean±SD** | **n** | **mean±SD** | **n** | **mean±SD** | **n** | **mean±SD** |
| 0 | 11 | 29.72 ± 23.50 | 3 | 63.91 ± 14.20 | 8 | 16.90 ± 6.53 | 11 | 29.72 ± 23.50 | 0 |  | 0 |  | 3 | 63.91 ± 14.20 |
| 7±1 | 4 | 17.37 ± 8.39 | 2 | 22.63 ± 7.04 | 2 | 12.11 ± 7.13 | 4 | 17.37 ± 8.39 | 0 |  | 0 |  | 2 | 22.63 ± 7.04 |
| 14±1 | 7 | 18.90 ± 9.73 | 4 | 15.03 ± 10.33 | 3 | 24.07 ± 7.33 | 6 | 19.45 ± 10.54 | 0 |  | 0 |  | 4 | 15.03 ± 10.33 |
| 21±1 | 5 | 20.76 ± 8.63 | 0 |  | 5 | 20.76 ± 8.63 | 3 | 21.54 ± 7.79 | 0 |  | 0 |  | 0 |  |
| 28±1 | 0 |  | 0 |  | 0 |  | 0 |  | 0 |  | 0 |  | 0 |  |
| 35±1 | 5 | 43.71 ± 23.72 | 0 |  | 5 | 43.71 ± 23.72 | 4 | 36.04 ± 18.91 | 1 | 74.40 | 0 |  | 0 |  |
| 37+ | 4 | 30.84 ± 9.19 | 0 |  | 4 | 30.84 ± 9.19 | 2 | 38.33 ± 3.40 | 2 | 23.35 ± 4.13 | 0 |  | 0 |  |

|  |  |  | **IBA1 perimeter-to-area ratio** | | | | | | | | | | | |
| --- | --- | --- | --- | --- | --- | --- | --- | --- | --- | --- | --- | --- | --- | --- |
|  | **Total** | | **CSF** | | **AM** | | **not treated** | | **viral treated** | | **PS (hCSF)** | | **PSA (hCSF)** | |
| **D** | **n** | **mean±SD** | **n** | **mean±SD** | **n** | **mean±SD** | **n** | **mean±SD** | **n** | **mean±SD** | **n** | **mean±SD** | **n** | **mean±SD** |
| 0 | 11 | 2.09 ± 0.37 | 3 | 2.11 ± 0.52 | 8 | 2.08 ± 0.35 | 11 | 2.09 ± 0.37 | 0 |  | 0 |  | 3 | 2.11 ± 0.52 |
| 7±1 | 4 | 3.61 ± 0.66 | 2 | 4.08 ± 0.02 | 2 | 3.14 ± 0.64 | 4 | 3.61 ± 0.66 | 0 |  | 0 |  | 2 | 4.08 ± 0.02 |
| 14±1 | 7 | 4.05 ± 0.70 | 4 | 4.46 ± 0.40 | 3 | 3.51 ± 0.68 | 6 | 4.08 ± 0.76 | 0 |  | 0 |  | 4 | 4.46 ± 0.40 |
| 21±1 | 5 | 3.79 ± 0.14 | 0 |  | 5 | 3.79 ± 0.14 | 3 | 3.77 ± 0.14 | 0 |  | 0 |  | 0 |  |
| 28±1 | 0 |  | 0 |  | 0 |  | 0 |  | 0 |  | 0 |  | 0 |  |
| 35±1 | 5 | 3.35 ± 0.28 | 0 |  | 5 | 3.35 ± 0.28 | 4 | 3.41 ± 0.27 | 1 | 3.09 | 0 |  | 0 |  |
| 37+ | 4 | 3.54 ± 0.47 | 0 |  | 4 | 3.54 ± 0.47 | 2 | 3.22 ± 0.48 | 2 | 3.86 ± 0.18 | 0 |  | 0 |  |

|  |  |  | **PV cell density (cells/mm2)** | | | | | | | | | | | |
| --- | --- | --- | --- | --- | --- | --- | --- | --- | --- | --- | --- | --- | --- | --- |
|  | **Total** | | **CSF** | | **AM** | | **not treated** | | **viral treated** | | **PS (hCSF)** | | **PSA (hCSF)** | |
| **D** | **n** | **mean±SD** | **n** | **mean±SD** | **n** | **mean±SD** | **n** | **mean±SD** | **n** | **mean±SD** | **n** | **mean±SD** | **n** | **mean±SD** |
| 0 | 14 | 32 ± 24 | 5 | 37 ± 30 | 9 | 29 ± 22 | 12 | 33 ± 26 | 0 |  | 2 | 27 ± 11 | 3 | 44 ± 40 |
| 7±1 | 9 | 12 ± 14 | 3 | 7 ± 11 | 6 | 14 ± 16 | 6 | 14 ± 16 | 0 |  | 3 | 7 ± 11 | 0 |  |
| 14±1 | 10 | 6 ± 10 | 4 | 1 ± 1 | 6 | 9 ± 12 | 3 | 4 ± 2 | 3 | 3 ± 5 | 2 | 0 ± 0 | 2 | 1 ± 1 |
| 21±1 | 6 | 6 ± 9 | 1 | 0 | 5 | 8 ± 10 | 4 | 10 ± 10 | 1 | 0 | 1 | 0 | 0 |  |
| 28±1 | 3 | 13 ± 9 | 0 |  | 3 | 13 ± 9 | 2 | 11 ± 11 | 1 | 17 | 0 |  | 0 |  |
| 35±1 | 9 | 7 ± 6 | 0 |  | 9 | 7 ± 6 | 5 | 6 ± 7 | 4 | 7 ± 4 | 0 |  | 0 |  |
| 37+ | 5 | 0 ± 0 | 0 |  | 5 | 0 ± 0 | 2 | 0 ± 0 | 3 | 0 ± 0 | 0 |  | 0 |  |

|  |  |  | **CR cell density (cells/mm2)** | | | | | | | | | | | |
| --- | --- | --- | --- | --- | --- | --- | --- | --- | --- | --- | --- | --- | --- | --- |
|  | **Total** | | **CSF** | | **AM** | | **not treated** | | **viral treated** | | **PS (hCSF)** | | **PSA (hCSF)** | |
| **D** | **n** | **mean±SD** | **n** | **mean±SD** | **n** | **mean±SD** | **n** | **mean±SD** | **n** | **mean±SD** | **n** | **mean±SD** | **n** | **mean±SD** |
| 0 | 11 | 70 ± 46 | 2 | 107 ± 73 | 9 | 62 ± 40 | 10 | 61 ± 38 | 0 |  | 1 | 158 | 1 | 56 |
| 7±1 | 7 | 34 ± 38 | 1 | 9 | 5 | 46 ± 39 | 5 | 46 ± 39 | 1 | 9 | 0 |  | 2 | 4 ± 6 |
| 14±1 | 9 | 26 ± 24 | 3 | 29 ± 20 | 6 | 25 ± 27 | 5 | 19 ± 12 | 2 | 25 ± 23 | 0 |  | 3 | 29 ± 20 |
| 21±1 | 6 | 12 ± 12 | 1 | 0 | 5 | 15 ± 12 | 5 | 14 ± 13 | 1 | 4 | 0 |  | 1 | 0 |
| 28±1 | 3 | 7 ± 4 | 0 |  | 3 | 7 ± 4 | 0 |  | 2 | 6 ± 5 | 0 |  | 0 |  |
| 35±1 | 5 | 21 ± 20 | 0 |  | 5 | 21 ± 20 | 4 | 23 ± 22 | 1 | 9 | 0 |  | 0 |  |
| 37+ | 3 | 14 ± 25 | 0 |  | 3 | 14 ± 25 | 1 | 0 | 2 | 21 ± 30 | 0 |  | 0 |  |

***Methods***

**Tissue preparation**

The distance of the obtained neocortical tissue from the tumour had been assessed by the neurosurgeon, based on magnetic resonance (MR) images, intraoperative pictures and occasionally defined by a navigational system. In case the obtained tissue was closer than 3 cm from the tumour, we considered it ‘close’, if the distance was more than 3 cm, we considered it ‘distant’.

The NMDG solution used for tissue transport and cutting^3^ contained (in mM): 93 N-Methyl-D-glucamine, 2.5 KCl, 1.2 NaH2PO4, 30 NaHCO3, 20 HEPES, 25 glucose, 5 ascorbic acid, 3 Na-pyruvate, 10 (MgSO4)7H2O and 0.5 (CaCl2)2H2O. The pH was adjusted with HCl in the range of 7.3-7.4, and the osmolarity was 300-305 mOsm. Although the dissection of the tissue block is in room air, all the tools are sanitised with 70% ethanol.

The slices were kept in ice-cold carbogenated NMDG until the end of the cutting to prevent tissue degradation. Then, slices were taken into a sterile hood, washed in warm culture medium (hCSF or AM) three times, by moving them through 3 plastic petri dishes, to prevent heat shock and to help recovery. The composition of the artificial culturing medium was the following (in mM): 8.4 g/L MEM Eagle medium, 20% heat-inactivated horse serum, 30 HEPES, 13 D-glucose, 15 NaHCO3, 1 ascorbic acid, 2 MgSO4, 1 CaCl2, 0.5 GlutaMAX-I, and 1 mg/L insulin, with a pH adjusted to 7.2–7.3, and osmolarity adjusted to 300–310 mOsm. The medium was sterile-filtered (pore size: 0.22 μm) and was stored at 4°C for up to two weeks (modified from ^3^). After the washing step, 1-3 slices were placed carefully on an insert (Millicell Standing Cell Culture Inserts, pore size 0.4 μm, diam. 30 mm, transparent PTFE membrane, hydrophilic, Merck Group, Burlington, Massachusetts, United States ) in a 6-well plate. The excess AM was removed from the top of the insert, and the wells were filled with 750 μl AM. The tissue slices were incubated for 1-4 hours with AM medium, then the culturing medium was changed by the same amount of AM supplemented with antibiotics (100unit/ml), either with penicillin and streptomycin (PS, Gibco, Thermo Fisher Scientific in Waltham, Massachusetts, USA)  or penicillin and streptomycin extended with amphotericinB (PSA, antibiotic+antimycotic solution, Sigma-Aldrich, Merck Group, Burlington, Massachusetts, United States) for one week to prevent infection. AM was changed every 3 days under sterile conditions in a laminar air flow with preheated AM, and the tissue cultures were kept at 35 °C, 5% CO2 and 100% humidity in the incubator.  After two weeks, a new batch of AM was prepared, and was introduced to the cultures stepwise in two feeding steps: the old and the new AM were mixed in 1:2, then in 2:1 ratio.

The clinical laboratory of the hospital collected the cerebrospinal fluid of patients undergoing lumbar puncture for their clinical investigation. The hCSF was collected and kept at 4°C for up to two weeks, then centrifuged. The supernatant was frozen in small aliquots in liquid nitrogene and kept at -80°C until use. The culture protocol for hCSF was the same as for AM.

**Recordings**

Electrophysiology

Both acute slices and organotypic cultures were transferred and maintained at 35–37°C in an interface chamber, perfused with a standard physiological solution containing (in mM) 124 NaCl, 26 NaHCO_3_, 3.5 KCl, 1 MgCl_2_, 1 CaCl_2_, and 10 D-glucose, equilibrated with 5% CO_2_ in 95% O_2_. We used a 24-contact (distance between contacts: 150 µm) laminar microelectrode^4,5^, and a custom-made voltage gradient amplifier of pass-band 0.01 Hz to 10 kHz. Signals were digitized with a 32 channel, 16-bit resolution analogue-to-digital converter (National Instruments, Austin TX, USA) at 20 kHz sampling rate, recorded with a home written routine in LabView8.6 (National Instruments, Austin TX, USA, RRID:SCR_014325). The linear 24 channel microelectrode was placed perpendicular to the pial surface, to record from every layer of the neocortex. Slices were mapped from one end to the other at every 300-400 µm. The pial surface could be identified throughout the whole culturing period by the naked eye. The microelectrode covered all layers of the neocortex. Usually, channels 1-12 were in the supragranular, channels 13-15 in the granular and channels 16-23 were in the infragranular layers. Channel positions were determined according to the thickness of the neocortex of the given tissue sample and corrected if necessary. The presence and the exact location of cellular activity (multiple unit or single unit activity) as well as that of population events were noted in case of every slice.

Wide-field imaging

Images were acquired using 2×2 spatial binning, resulting in a field of view of approximately 13.3×13.3 mm and a spatial resolution of 13.0 µm/pixel. Acquisition parameters varied by patient cohort: recordings for patients HOC13 and HOC15 were captured at 10 frames per second (fps) with a 10 ms exposure time, while recordings for patient HOC21 were acquired at 30 fps with a 33 ms exposure time.

**Data analysis**

Electrophysiology

Detection of the SPA was performed on LFPg recordings after a Hamming window spatial smoothing and a band-pass filtering between 1 and 30 Hz (zero phase shift, 48 dB/octave). Note that SPA is the totality of ‘SPA events’ occurring in a given recording (on average 193±235 SPA events/recording). SPA events were visible usually on 5 to 16 channels, and SPA events larger than two times the standard deviation of the basal activity were detected and included in the analysis. The largest amplitude LFPg peak of the events was chosen as time zero for averaging. The location and the recurrence frequency of the SPA events were determined in each file during the whole recording session. Current source density (CSD, an estimate of population transmembrane currents) and multiple unit activity (MUA, an estimate of population neuronal firing) were calculated from the LFPg using standard techniques^6^. LFPg, CSD and MUA were averaged from -300 to +300 ms from the peak of the events determined as described above. Baseline correction (-250 to -100 ms) was applied to averaged LFPg, CSD and MUA.

For neuron clustering, broad-band electrophysiological recordings were bandpass-filtered (500–5000 Hz, 4th-order zero-phase Butterworth filter) to isolate single-unit spiking activity. Initial spike detection and cluster assignment were performed with Kilosort4^7^, curated with phy (Cortex Lab), yielding spike times, spike cluster identities and metadata. Further manual curation was carried out using a custom Python-based spike viewer (availability upon request). Excitatory principal cells (PC) and inhibitory interneurons (IN) were separated based on their AP width and firing characteristics. Based on their firing characteristics on the autocorrelogram, PCs were further divided into intrinsically bursting (IB) and regular spiking (RS) principal cells^2^. The location of the single cells was determined in each recording, as described above (SPA location). For each unit, spike waveforms were extracted from the target channel in a ±2 ms window around each detected spike time and averaged to assess waveform quality. Units were manually labelled as "good" or "noise" based on waveform shape, firing rate, and inter-spike interval distributions visible in the viewer. Individual spikes could be added by left-clicking at the location of a detected peak (resolved to the absolute, positive, or negative extremum within a ±1.5 ms neighbourhood), or removed by right-clicking within a ±0.5 ms window around an existing spike. Contiguous suprathreshold intervals on the bandpass-filtered trace were used to detect population events for a designated channel, retaining one peak per interval to avoid double-counting. Clusters could be merged, split by an interactively set amplitude threshold, or reassigned across units. All modifications—including spike additions, deletions, and cluster remodelling – were saved back for downstream analysis. Only neurons with a clear refractory period of at least 2 ms were included. We were aware that the AP waveform slightly changes during burst activity and checked all presumptive APs accordingly. AP waveform analysis was performed in Python 3.14, on averaged APs made from wide-band recordings.

Single-unit recordings were classified at two levels of granularity. At the *fine* level, units were assigned to one of three electrophysiological subtypes: regularly spiking principal cells (RS-PC), intrinsically bursting principal cells (IB-PC), and interneurons (IN), based on waveform characteristics and firing properties. At the *coarse* level, RS-PC and IB-PC units were pooled into a single principal cell (PC) category and compared against interneurons.

Wide-field imaging

The following image processing steps were applied. Rolling-ball background subtraction was applied to this mean image to correct for uneven illumination. Adaptive global thresholding (triangle method) generated a binary tissue mask, which was used for morphological closing, hole-filling, and extraction of the largest connected component subsequently refined to isolate the tissue slice. The fluorescence movie was then restricted to this segmented mask, collapsing the spatial dimension in each frame to yield a single mean fluorescence trace representing whole-slice activity dynamics. To remove slow signal drift, a third-degree polynomial baseline was fit and subtracted from the trace.

**Histology**

Immunostaining protocol

After fixation, acute and cultured slices were recut to 60 µm thick sections and thoroughly washed with 0.1M PB. Usually, one to several sections derived from the same slice were selected for the different stainings (see below). Sections were transferred to 0.1M Tris-buffered saline (TBS, pH:7.4), then endogenous peroxidase was blocked by 1% H_2_O_2_ in TBS for 10 minutes. TBS was used for all the washes (3x10 min between each serum) and for the dilution of the antisera. Non-specific immunostaining was blocked by 2% normal goat serum and 2% normal horse serum for one hour. Sections were processed for immunostaining against the neuronal cell body marker NeuN antibody (1:2000, EMD Millipore, Billerica, MA, USA, RRID:AB_2298772), and the astroglial cell marker glial fibrillary acidic protein antibody (GFAP, 1:2000, EMD Millipore, Billerica, MA, USA, RRID:AB_94844), the microglia marker IBA1 (1:2000, Invitrogen, Waltham, Massachusetts, USA, RRID: AB_2544912), the inhibitory cell markers parvalbumin (PV, 1:7000, Swant, Bellinzona, Switzerland, RRID:AB_10000343), calretinin (CR, 1:2000, Swant, Bellinzona, Switzerland, RRID:AB_10000320) or the green fluorescent protein for viral expression detection (GFP, Nacalai Tesque, Kyoto, Japan, RRID:AB_10013361). The IBA1 antibody was a rabbit polyclonal, the GFP antibody was a rat monoclonal, all the other antibodies were mouse monoclonal antibodies. Their specificity was tested by the manufacturer. The primary antibodies were applied for two days at 4°C. For visualization of the immunopositive elements biotinylated anti-rabbit (in case of IBA1), biotinylated anti-rat (for GFP) or anti-mouse (in case of NeuN, GFAP and PV) IgG (1:250, Vector, Burlingame, CA, USA) was applied as secondary antibody (2 hours), followed by avidin-biotinylated horseradish peroxidase complex (ABC, 1:250, 1.5 hours, Vector). The immunoperoxidase reaction was developed by 3,3’-diaminobenzidine tetrahydrochloride dissolved in 0.05M Tris buffer (pH:7.6), as a chromogen. Sections were mounted on slides from chrome-gelatine solution (solved in 0.1M PB), air dried and mounted in DePeX (Electron Microscopy Sciences, Hatfield, PA, USA).

**Data acquisition**

Sections stained against NeuN, GFAP, IBA-1, PV and CR were scanned with a Pannoramic MIDI III scanner (3D Histech Ltd. Budapest, Hungary, RRID:SCR_027467). In each case, every slice was scanned entirely at a magnification of 20x.

The density of NeuN-positive neurons and the perimeter-to-area ratio of IBA1-positive microglial cells were determined with the aid of the QuPath v0.6.0. software^8,9^ and StarDist extension with the pre-trained model 2D_paper_dsb2018. The density of CR-positive and PV-positive interneurons was determined in a semi-automated manner with the help of the above-mentioned extension combined with the built-in cell detection of QuPath. In the case of the NeuN, PV and CR staining, cell bodies located in the entire section were counted, and the area of the scanned section was also determined with pixel-classification. In the case of the IBA1 staining, a 1600x1200-pixel-sized rectangle was placed on a representative area containing at least 50 cell bodies to calculate the perimeter-to-area ratio of the detected microglial cells.

GFAP-positive coverage, and IBA1-positive coverages were measured with pixel classification trained on multiple representative areas in QuPath (for details see below).

If more than one section was stained and scanned from the same slice with the same staining, all sections were used for the quantitative analysis, weighted with the area (and the standard deviation was calculated between the sections). Together with the acute slices, a total of n=179 slices were processed, n=134 slices used for NeuN, n=124 slices for GFAP, n=51 slices for IBA1, n=86 slices for PV and n=54 slices for CR cell counting. Virally treated slices were treated separately, as virus injection can induce different changes compared to normal culturing.

**Detection for quantitative analyses**

Settings in the QuPath program to detect **NeuN-positive** cells on n=134 sections:

- Model: dsb2018_heavy_augment.pb; detection threshold 0.6; pixel size 0.5 µm.
- Preprocessing: H-DAB deconvolution, extract channels 0 and 1, sum, median(1), divide by 2, normalize 1–99 percentiles.
- Valid cell filter: Area 15–800 µm²; Solidity ≥ 0.6; Circularity ≥ 0.5; Aspect ratio ≤ 2.6; DAB Std.Dev. ≥ 0.04; Hem Std.Dev. ≥ 0.04.
- Dendrite removal: Area < 40 µm², remove if buffered by 15 µm and intersects a neighbour with Area > 100 µm².

Settings for pixel classification trained on 7 sections stained **against GFAP**, then applied on n=124 sections:

- Classifier: OpenCVPixelClassifier with RTrees (50 trees), classification task with two outputs (Glia, Tissue).
- Features: Multiscale Gaussian + Laplacian on RGB at σ = 0.5, 1.0, 2.0, 4.0 (24 total features), with identity normalisation.
- Inputs: RGB tiles 512×512, pixel size ~0.217391 µm/px (smallest available), no stain deconvolution, no extra preprocessing beyond the multiscale filters.

Settings for pixel classification trained on 4 sections stained against **anti-IBA1**, then applied on n=51 sections:

- Classifier: RTrees inside OpenCVPixelClassifier, 50 trees, classification task with classes Other and PC.
- Features: GAUSSIAN and LAPLACIAN on RGB at σ=1.0, 6 features total, identity normalisation.
- Inputs: RGB tiles 512×512, pixel size ~0.217391 µm/px (smallest available), no stain deconvolution or extra preprocessing beyond the multiscale filters.

Settings to calculate perimeter-to-area ratio on a region of interest in **IBA1 stained** sections containing at least 50 somas, if possible, applied on n=51 sections:

- Model: dsb2018_heavy_augment.pb; detection threshold 0.4; pixel size 0.3 µm
- Preprocessing: H-DAB deconvolution, extract channels 0 and 1, sum, median(1), Gaussian blur (σ = 1.0), multiply by 1.5, normalise 0.5–99.5 percentiles.
- Valid cell filter: Nucleus area 20–800 µm²; Circularity 0.55–0.99; Solidity ≥ 0.6; DAB positivity (nucleus or cell DAB mean > 0.09); nuclear hematoxylin mean > 0.12.
- Probability filter: Detection probability ≥ 0.55.
- Classification: IBA1+ Activated vs Resting (Cell DAB mean > 0.3 threshold) crossed with Large vs Normal nucleus (Nucleus area ≥ 100 µm²), giving 4 final classes.
- The areas and perimeters of the 50 detections with the highest detection probability were saved for further analysis.
- Calculation: $x=\frac{area}{perimeter}\approx\frac{\pi r^{2}}{2\pi r}\Rightarrow$the bigger the x, the more activated the microglia

**Statistical tests used for the histological analysis:**

| Normality | 2 groups, paired | 2 groups, unpaired | Multiple groups, paired | Multiple groups, unpaired |
| --- | --- | --- | --- | --- |
| Parametric | Paired t-test | Unpaired t-test with Welch’s correction | ANOVA Greenhouse- Geisser correction  +multiple comparison | ANOVA + multiple comparison |
| Non-parametric | Wilcoxon signed rank test | Mann-Whitney U test | Friedman test + multiple comparison | Kruskal-Wallis ANOVA + multiple comparison |

**Statistics for the electrophysiology data**

The following electrophysiological metrics were included in all comparative and correlative analyses: firing rate (Hz), maximum firing rate over the most active 10-second window (Hz), burstiness index 1 (%), burstiness index 2, based on inter-spike intervals shorter than 20 ms (%), spike half-width measured at main, and if present, auxiliary peaks (half-width 1 and half-width 2, in ms), inter-spike interval coefficient of variation (ISI CV). For each metric, descriptive statistics (mean ± standard deviation and sample size n) were calculated separately for each cell type (fine and coarse), each patient, each day in vitro (DIV), and for pooled groups (all principal cells, all cells, D0 versus D>0).

Spearman rank correlation coefficients (ρ) were computed between all pairs of electrophysiological metrics within each cell type, to characterise co-variation of firing properties independently of distributional assumptions. Correlation matrices were visualised as heatmaps with significance markers overlaid (Bonferroni-corrected within each matrix). To assess developmental trajectories, Spearman correlations were additionally computed between DIV and each metric within each cell type. Median values per DIV were plotted as trend lines overlaid on scatter distributions to aid visual interpretation.
